# scPrisma: inference, filtering and enhancement of periodic signals in single-cell data using spectral template matching

**DOI:** 10.1101/2022.06.07.493867

**Authors:** Jonathan Karin, Yonathan Bornfeld, Mor Nitzan

## Abstract

Single-cell RNA-sequencing has been instrumental in uncovering cellular spatiotemporal context. This task is however challenging due to technical and biological noise, and as the cells simultaneously encode multiple, potentially cross-interfering, biological signals. Here we propose scPrisma, a spectral computational framework that utilizes topological priors to decouple, enhance, and filter different classes of biological processes in single-cell data, such as periodic and linear signals. We demonstrate scPrisma’s use across diverse biological systems and tasks, including analysis and manipulation of the cell cycle in HeLa cells, circadian rhythm and spatial zonation in liver lobules, diurnal cycle in Chlamydomonas, and circadian rhythm in the suprachiasmatic nucleus in the brain. We further show how scPrisma can be used to distinguish mixed cellular populations by specific characteristics such as cell type, and uncover regulatory networks and cell-cell interactions specific to predefined biological signals, such as the circadian rhythm. We show scPrisma’s flexibility in utilizing diverse prior knowledge, and inference of topologically-informative genes. scPrisma can be used both as a stand-alone workflow for signal analysis, and, as it does not embed the data to lower dimensions, as a prior step for downstream single-cell analysis.

## 1 Introduction

In recent years, progress in single-cell RNA sequencing (scRNA-seq) that contains information about the gene expression profiles of multitude of cells across tissues, has led to significant improvement in our understanding of a variety of intracellular and intercellular processes [47]. Computational advances developed over the past few years have pushed forward the interpretation of this data to extract information about heterogeneity of cell types and states and their collective structure and behavior, including spatial context [31], gene regulatory patterns [18], cell type [35], and temporal processes such as lineage [12] and cell cycle [25]. Since scRNA-seq data contains multiple biological signals, it can be challenging to uncover a particular underlying signal, and in many cases, prior information about the signal in question is needed [1, 25]. Recently, we have shown how topological priors about the hierarchical structure of lineage and differentiation relationships in single-cell data can be leveraged for their identification using spectral approaches, as they exhibit power-law signature in the covariance eigenvalue distribution [30]. Relying on global topological features (e.g. hierarchical characteristics of a signal) has the advantage of being a more robust and generalizable approach than relying on specific features such as marker gene information [1, 25], as these are many times unique to different biological systems and processes and can be challenging to infer for new systems. Here, we show how topological priors can be used beyond signal identification. We use priors about the periodicity and linearity of diverse biological processes to either enhance or filter them out from single-cell data using a spectral projection approach.

Biological processes that are inherently periodic are abundant and play important roles in diverse contexts, such as the cell cycle and circadian rhythm. Characteristics of the cell cycle have been studied extensively in the last decades [21], and recent studies of its gene expression dynamics using scRNA-seq have been mostly focused on one of two aspects; First, studying the dynamics and the topology of the cell cycle in gene expression space [41] and its coupling to other biological processes [20]. Second, since the cell cycle is one of the major determinants of gene expression variance in many biological systems [22, 28], filtering out its influence can improve downstream analysis related to additional processes or information encoded by the cells, such as cell type representation[1]. The circadian rhythm and its representation in various tissues has also been the focus of extensive scRNA-seq studies in recent years [7, 11, 26]. A disruption of the circadian rhythm can lead to adverse health effects, such as metabolic disorders and liver disease[43], cancer [40], or heart disease[38], and thus, better characterization of its properties can provide a key to both basic understanding of such disorders as well as lead to new therapeutic strategies[38].

Multiple computational methods aim to infer periodic signals from single-cell data, with a particular focus to extract information related to the cell cycle or to remove its effect[1, 3, 24, 25]. However, the majority of these computational methods are heavily based on cell cycle marker gene information, including ccRemover (a correlation-based approach[1]), Seurat (which extracts the cell-cycle factor from residual gene expression based on the expression of cell cycle markers), and reCAT[25] (based on an approximation of the traveling salesman problem). Their reliance on specific marker gene information makes these approaches difficult to generalize across systems and across additional periodic signals beyond the cell cycle. On the other hand, Cyclum[24], which does not rely on marker gene information, is an auto-encoder based approach that optimizes a circular embedding for single-cell data to infer and remove cell cycle effects. However, fitting the data to a 1-dimensional circle, as is done in Cyclum, does not generally capture the variability of cyclic biological processes and risks being too restrictive, and, being a black-box method that does not readily integrate additional prior knowledge (e.g. low-resolution temporal information) reduces the flexibility of the method, as such additional data may be necessary in cases of weak cyclic signals.

In this paper, we develop scPrisma, a generalized spectral framework (Figure 1) for the reconstruction, enhancement, and filtering of cyclic signals, as well as inference of informative cyclic genes, and is further extended to linear signals. We benchmark our approach and demonstrate its performance over simulated data and four scRNA-seq datasets. Specifically, we show how the cell cycle can be revealed or filtered in a population of HeLa cells, how circadian rhythm and spatial zonation can be decoupled in liver lobules, how differences in Chlamydomonas that were grown in different environments can be emphasized by filtering their diurnal cycle signal, and how the signature of the circadian rhythm can be revealed in multiple cell types in the suprachiasmatic nucleus (SCN) in the brain, the master circadian pacemaker in mammals. In addition, we show how the enhancement of specific biological signals using scPrisma allows us to better distinguish distinct cellular subtypes of SCN neurons following temporal filtering, and uncover signal-related gene regulatory networks and cell-cell interactions following enhancement of the circadian rhythm signal.

**Figure 1:**
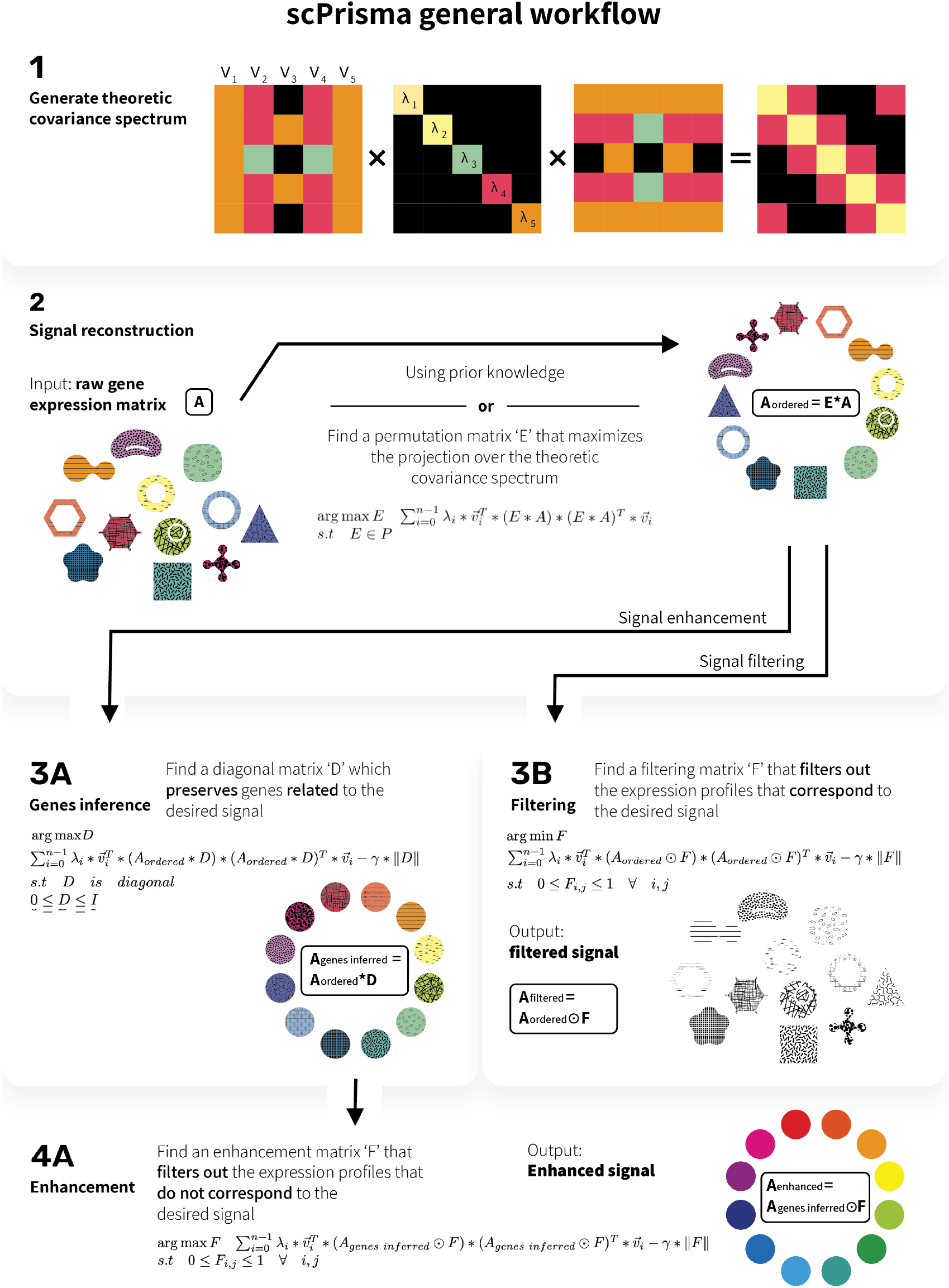
General workflow of scPrisma

scPrisma is versatile and enables reconstruction, gene inference, filtering, and enhancement of the underlying cyclic or linear signals of interest, without low-dimensional embedding, which renders the results useful for diverse types of downstream analyses. Furthermore, it is flexible as it enables integration of diverse types of prior knowledge (such as low-resolution temporal ordering, but does not rely on it and can be used for de novo analyses.

## 2 Results

### 2.1 scPrisma

We developed scPrisma, a spectral analysis framework that uses topological priors over underlying signals in single-cell data, to allow for their inference, enhancement, and filtering. The core of scPrisma utilizes spectral template matching between the spectrum (the eigendecomposition of the covariance matrix) of a set of single-cell data (e.g. scRNA-seq), and the expected analytical spectrum of a structure or process we aim to enhance or filter. To analyze a theoretical covariance spectrum (by analyzing its eigenvalues and eigenvectors), we need a reference model. Focusing first on cyclic signals, we developed a simple toy model of periodic biological signals (as described in Methods 4.1). The covariance matrix of the gene expression matrix of this model is a circulant matrix of a special form that depends on the model parameters (Methods 1). Circulant matrices have closed form formula for their eigenvectors and eigenvalues [37] (Figure 1.1), which we use in order to estimate the ordering of cells along the cyclic topology. This is done by optimizing for a permutation matrix that maximizes the projection of the data over the theoretical spectrum (Methods 4.5; Figure 1.2). As an input for the remainder of scPrisma’s workflow, cellular ordering can also be informed by prior knowledge of low resolution pseudotime. Based on the reconstructed ordering, scPrisma infers topologically informative genes, as the set of genes that maximizes the projection over the theoretical spectrum (Methods 4.6; Figure 1.3A). scPrisma can then either enhance or filter out the signal related to the cyclic process by filtering out gene expression entries that do not maximize or minimize, respectively, the projection over the theoretical spectrum (Methods 4.7; Figure 1.4A, Figure 1.3B).

### 2.2 scPrisma identifies and filters periodic signals

As a starting point, we assess the performance of scPrisma on simulated data and show its ability to reconstruct underlying cyclic signals, infer cyclic genes, as well as filter or enhance cyclic signals. We first simulated noiseless gene expression matrices encoding a cyclic signal (Supplementary methods A.1). scPrisma successfully captures the cyclic signal (Figure 2A), and the reconstruction is robust to noise, as the quality of reconstruction gradually decreases with decreasing SNR (signal to noise ratio, 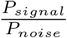, where we use Frobenius norm as power), and deteriorates only at high noise levels (*SNR* < 0.5) (Figure 2B; Supplementary A.2; Supplementary Figure 1).

**Figure 2:**
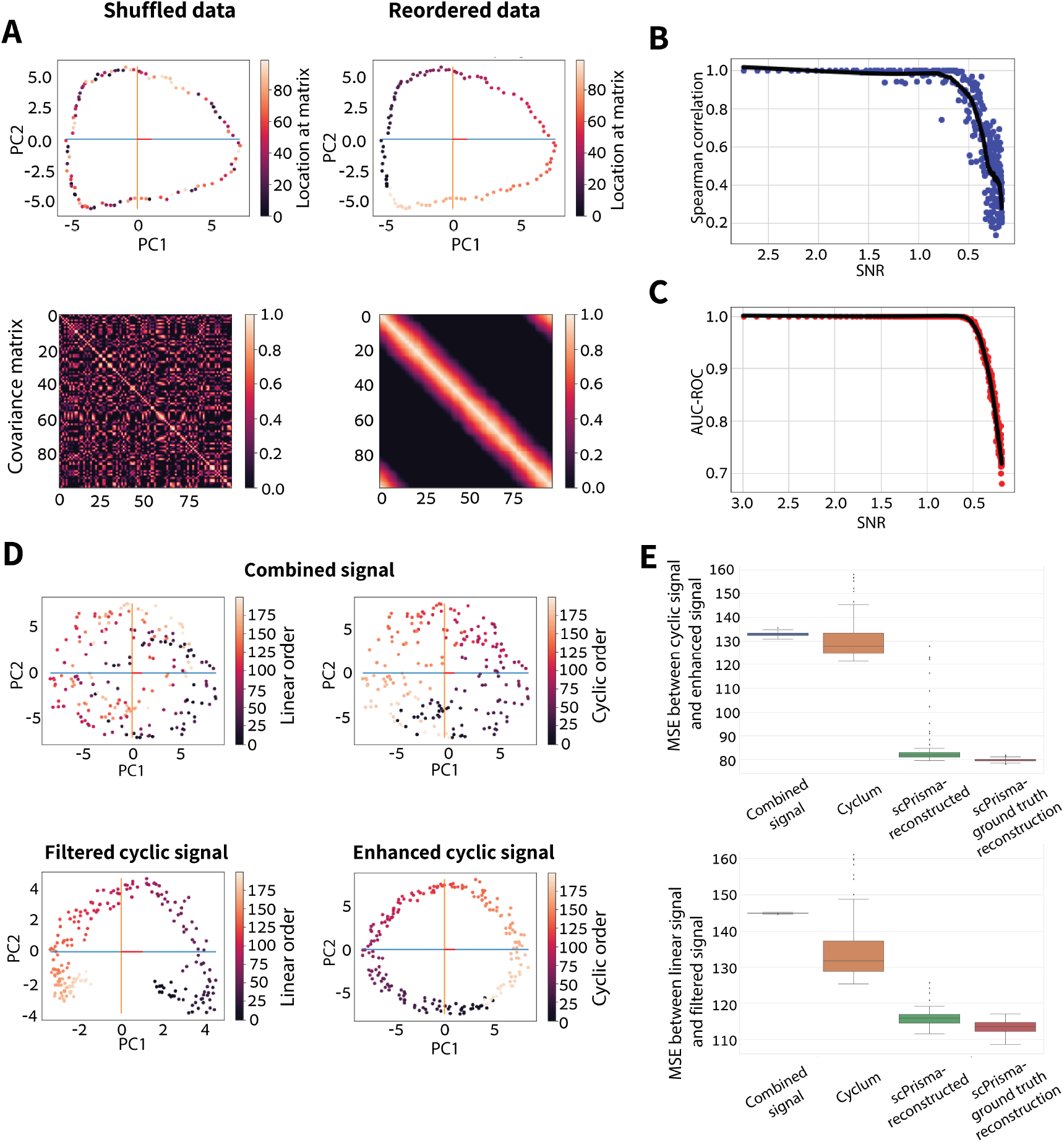
scPrisma identifies and filters periodic signals in simulated data. (A) PCA representation and gene expression covariance matrix before and after applying the reconstruction algorithm. This was done over a simulated gene expression matrix (100 cells, 500 genes) encoding a cyclic signal according to the spatial model (Supplementary A.1). The covariance matrix, after applying the reconstruction algorithm, is circulant. (B) Spearman correlation between the ground-truth permutation and the predicted permutation as a function of SNR over 300 simulations. A cyclic signal was simulated similarly to (A), and Gaussian noise was added with varying variance. (C) AUC-ROC of informative genes inference task as a function of SNR over 300 simulations, each consisting of a cyclic signal and a lineage signal (256 cells, 250 genes) that were concatenated to form a combined gene expression matrix (256 cells, 500 genes), supplemented by additive Gaussian noise. (D) Filtering and enhancement of cyclic signals in combined simulated data. The combined simulation consists of the sum of linear and cyclic signals (200 cells, 500 genes each), and a Gaussian noise matrix (variance = 0.1). (E) Boxplots comparing the filtering and enhancement algorithms of scPrisma with Cyclum over 100 simulations of the combined signal as described in (D). For the box plots, the center line is the median, box limits are the 0.25 and 0.75 quantiles, vertical lines extend from the top of the box to indicate the maximum value, and from the bottom to indicate the minimum value.

Using scPrisma, we can also correctly classify the subset of cyclic genes and distinguish them from genes expressing Gaussian noise (Supplementary A.2). Further, in a more challenging setting, when entangling the cyclic signal with a hierarchical signal (that can capture lineage relationships within the data, following the model suggested in [30]; Supplementary A.2), our approach can still identify the subset of cyclic genes, in a way that is robust to increasing noise (Figure 2C).

Next, to demonstrate the capabilities of our approach for either filtering or enhancing underlying cyclic signals in the data, we simulated noisy gene expression matrices with embedded cyclic as well as linear signals (Methods 4.2). scPrisma can both enhance, as well as filter the cyclic signal, and thus, either weaken or retain and effectively enhance the linear signal in the data, respectively. For this task we benchmarked scPrisma relative to Cyclum[24], which supports the inference and filtering of circular components in scRNA-seq data. scPrisma robustly outperforms Cyclum in both enhancement of the cyclic signal (mean MSE between the original gene expression matrix and the cyclic signal reduces from 133.10 to 84.45 with scPrisma, compared to 130.87 with Cyclum, and 79.99 with ground-truth reconstruction, Figure 2E), as well as filtering of the cyclic signal, thus revealing the underlying linear signal in the data (mean MSE between the original gene expression matrix and the linear signal reduces from 144.90 to 115.91 with scPrisma, compared to 134.57 with Cyclum, and 113.70 with ground-truth reconstruction, Figure 2E). Furthermore, when simulating a cyclic signal with varying amount of Gaussian noise, we found a substantial improvement in recovering the underlying cyclic signal following its spectral enhancement (Supplementary Figure 2C). Finally, when simulating single-cell data combining linear and hierarchical signals, and using scPrisma’s linear spectral analysis (Methods 4.2), we find that the inference of genes associated with the linear process is robust and degrades only at high noise levels (*SNR* < 0.5) (Supplementary Figure 2E,F).

### 2.3 scPrisma manipulates the cell cycle signal in HeLa cells

We next tested our approach on a scRNA-seq dataset of HeLa cells (Figure 3), unsynchronized across the cell cycle [41]. To assess the results we used a list of 300 genes, classified and clustered according to cell cycle phases (G1.S, S, G2, G2.M, M.G1)[41], where for each phase, the corresponding genes were summed and normalized (Supplementary A.3), and their circular mean and variance were calculated [17] (Supplementary A.3). For an ordered reconstructed signal, the ordering of the circular means should correspond to the cell cycle phases and the circular variance should be less than 1, while for randomly ordered data, the circular variance is expected to be close to 1 (corresponding to a uniform signal along the cycle). Following standard preprocessing of the HeLa single-cell data (Methods 4.3, Supplementary A.3), the distributions of all phases are found to be nearly uniform (Figure 3B, mean circular variance=0.994). However, after cyclic ordering by scPrisma (Methods 4.5), different phases of the cell cycle become clearly separated (Figure 3B, mean circular variance=0.850) and peak progressively according to the correct phase ordering (the circular mean in radians: G1.S: 1.407, S: 0.323, G2: −0.531, G2.M: −0.721, M.G1: −1.154.).

**Figure 3:**
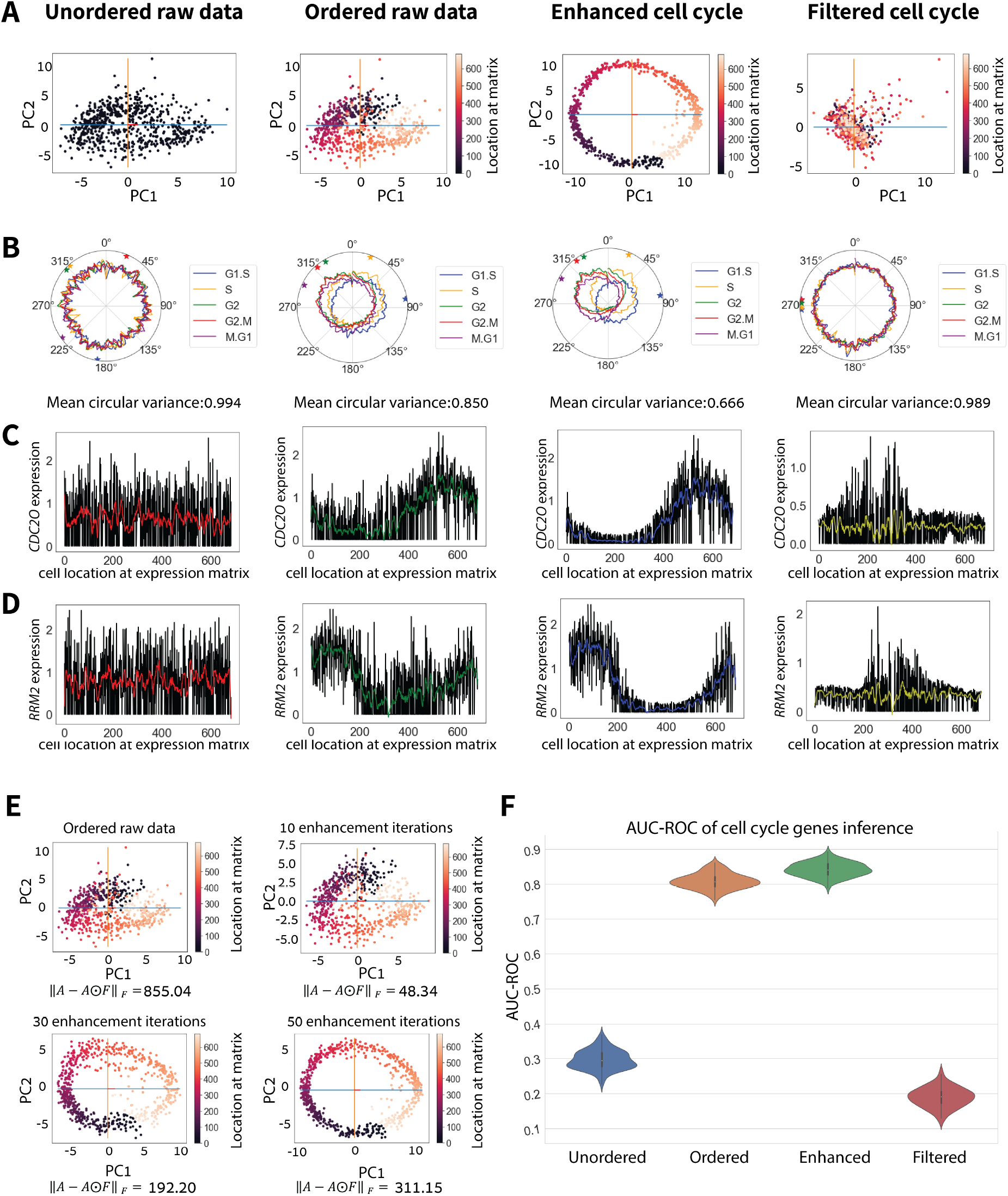
scPrisma manipulates the cell cycle signal in HeLa cells. (A) PCA representation of cells in the unordered and ordered raw gene expression data, as well as data following spectral cyclic enhancement and filtering. The cells are colored according to the corresponding rows in the gene expression matrix. (B) Smoothed polar plot of the normalized sum of the gene sets corresponding to different cell cycle phases[41]. The circular mean of each phase is marked by a correspondingly colored star. (C,D) Expression of *CDC20* (which peaks in ‘M’ phase [39]) and *RRM2* (which peaks in ‘S’ phase [39]) as a function of cellular location in the gene expression matrix. (E) PCA representation of cells along iterations of the cyclic enhancement algorithm. (F) Violin plot of AUC scores for 50 random gene subsets inferred by scPrisma as cyclic genes based on unordered, ordered, enhanced and filtered data.

To further enhance the cell cycle signal in the data, we employed scPrisma’s cyclic enhancement algorithm. Iterations of our algorithm gradually reveal a cyclic signal, which is apparent after filtering less than 7% of the total signal, and is clearly revealed in the reduced 2-dimensional principal component space after removing 22% of the total signal (Figure 3E). In addition, the reconstructed angular ordering is correlated with the cell cycle phases, and the circular variance of phase-corresponding marker genes decreases substantially (Figure 3B, mean circular variance=0.666). Further, genes associated with different cell cycle phases peak progressively in their expected order along the enhanced cycle (Methods 4.7, Figure 3C,D).

Next, we approached the reverse challenge: to filter out the cell cycle signal from the HeLa cells data. After applying our cyclic filtering algorithm on the ordered data, the separation between different cell cycle phases and their corresponding progressive peaks are lost, (Figure 3B, mean circular variance=0.989).

Finally, we identify genes related to the cell cycle using the genes inference algorithm (Methods 4.6, Supplementary A.3). When using as input both randomly-selected subsets of cell cycle related genes[41] and subsets of genes unrelated to the cell cycle, we find a substantial improvement in our ability to identify cell cycle related genes after reordering the cells (mean AUC = 0.808) relative to the original data (mean AUC = 0.297). Furthermore, relative to the ordered data, identifying cell cycle related genes is further improved following spectral cyclic enhancement (AUC mean 0.841), and is diminished following cyclic filtering (AUC mean 0.189) (Figure 3F).

### 2.4 scPrisma Disentangles spatial and temporal signals in liver lobules

We next attempt to dissect spatiotemporal signals encoded by cells via a scRNA-seq dataset which captures gene expression variation of hepatocytes in the mammalian liver across both space (spatial zonation across the periportal to pericentral axis) and time (temporal variation across the circadian rhythm) [7]. Similarly to the cell cycle, the circadian rhythm is also expected to exhibit a cyclic structure in gene expression space. In this experiment[7], liver cells were sequenced at four equally-spaced time points along the day, and therefore, we will use this prior knowledge for ordering the cells in a cycle according to their low-resolution sampling time. In this setting, we are still missing information about temporally informative genes, and spatiotemporal information is still entangled (e.g. *Pck1* varies informatively across both space and time of day[7]). Therefore, we leverage scPrisma to disentangle this single-cell data, and we show clear enhancement, as well as filtering of the circadian rhythm, relative to raw data (K=4 Adjusted Rand Score (ARI) of KMeans=0.96; 0.013; 0.11, respectively; Figure 4A,E; Methods). Next, we focus on the analysis of individual genes. *Pck1* is a rhythmic gene that is known to be highly expressed at two out of the four measured timepoints, ZT06 and ZT12 [7], and indeed, following spectral cyclic enhancement, its resulting expression in ZT00 and ZT18 is diminished (Figure 4C). Conversely, cyclic filtering resulted in a nearly constant expression of *Pck1* across the four time points along the circadian rhythm. Spatially, *Pck1* is known to be periportally zonated (its expression is monotonically increasing along the pericentral to periportal axis[15]) (Figure 4C). As expected, spectral cyclic enhancement flattens *Pck1* expression across the lobule layers, while cyclic filtering retains the variation across space. Similar behavior can be observed for additional spatiotemporally informative genes (Figure 4D, Supplementary Figure 3).

**Figure 4:**
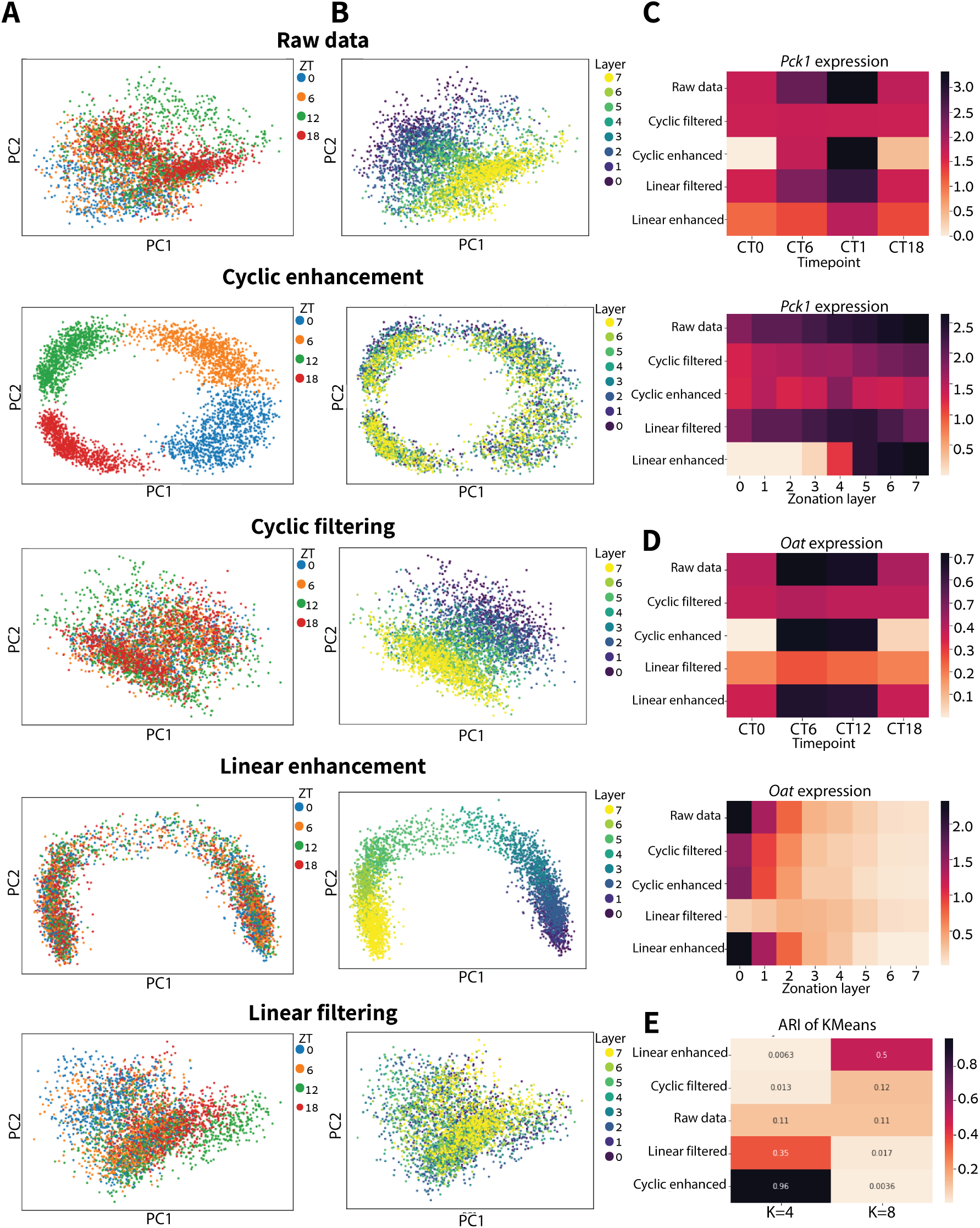
Disentanglement of spatial and temporal signals in liver lobules. (A,B) PCA representation of raw single-cell data, as well as following spectral cyclic enhancement, cyclic filtering, linear enhancement, and linear filtering. The cells are colored either according to their associated time points (‘ZT’, sampled at four equally-spaced timepoints along the circadian rhythm) (A), or by their respective spatial location (‘Layer’, according to the zonation analysis done by[7]) (B). (C,D) Heatmaps of the expression of *Pck1* (C) and *Oat* (D) following scPrisma analysis as a function of the sampling time and zonation layer. (E) Adjusted Rand Score (ARI) of KMeans clustering with K=4 (corresponding to 4 underlying time points) and K=8 (corresponding to 8 underlying zonation layers) after applying each of scPrisma’s spectral algorithms.

In a complementary manner, we next show that spectral analysis focusing on the characteristics of linear signals can be used to filter the spatial linear signal in the collective gene expression of hepatocytes (Methods 4.7). Indeed, following spectral filtering by scPrisma, the cyclic circadian signal is clearly revealed (Figure 4A) and the linear zonation signal is blurred out relative to the raw data (Figure 4B,E; K=8 ARI of KMeans=0.017; 0.11, respectively). Focusing again on *Pck1* expression, linear spectral enhancement indeed retains only the expression around the portal vein and reduces the variance across different timepoints (Figure 4C,E; K=8 ARI of KMeans=0.5), while linear filtering reduces the zonation variance yet keeps the variance between different timepoints (Figure 4C).

Finally, we evaluated the dominant structure in the data following scPrisma analysis. As predicted, following spectral enhancement, the data clusters according to the enhanced signal, while following spectral filtering, the data clusters according to the unfiltered signal; for example, filtering the cyclic signal leads to better identification and clustering of the spatial zonation signal (Figure 4E).

### 2.5 scPrisma detects and filters the diurnal cycle in Chlamydomonas

To demonstrate the use of scPrisma for more complex systems with diverse prior knowledge, we next turn our attention to scRNA-seq data collected for Chlamydomonas (green algae), grown under two contrasting conditions, iron replete (Fe+) and iron deficient (Fe-) [27]. In both conditions, a signal in the gene expression of the Chlamydomonas that reflects the 24 hours diurnal cycle was previously detected [27]. To evaluate progression of cells along the diurnal cycle, we used marker genes corresponding to different cycle phases obtained from bulk RNA-sequencing [44] (Supplementary A.5). Using scPrisma’s cyclic enhancement resulted in robust reconstruction of the diurnal cycle for each of the two conditions (Methods 4.5; Figure 5A for FE+ condition; Supplementary Figure 4B for FE-condition). Further, concatenating the enhanced cyclic signal of both experiments resulted in a reconstructed synchronized diurnal cycle (Supplementary Figure 3A), and therefore, filtering out this joint signal is expected to enhance the underlying divergence between the experiments. Therefore, we next focused on enhancing the biological differences between the FE- and FE+ conditions by spectrally filtering their shared diurnal cycle (Figures 5, Supplementary Figure 4). As expected, cyclic filtering increased the differences between the clusters of FE- and FE+ associated cells (Silhouette score before/after filtering = 0.088/0.136;, Figure 5B). scPrisma outperformed the results of available, alternative cyclic filtering methods (Supplementary Methods A.7; Figure 5B), including ccRemover[1], Seurat[3] and Cyclum[24] (Silhouette scores = 9.815e-06, 9.868e-04, and 0.052, respectively). Methods that exploit the cyclic nature of the diurnal cycle, such as scPrisma as well as Cyclum, perform better than methods that heavily rely on cyclic marker genes (such as Seurat and ccRemover).

**Figure 5:**
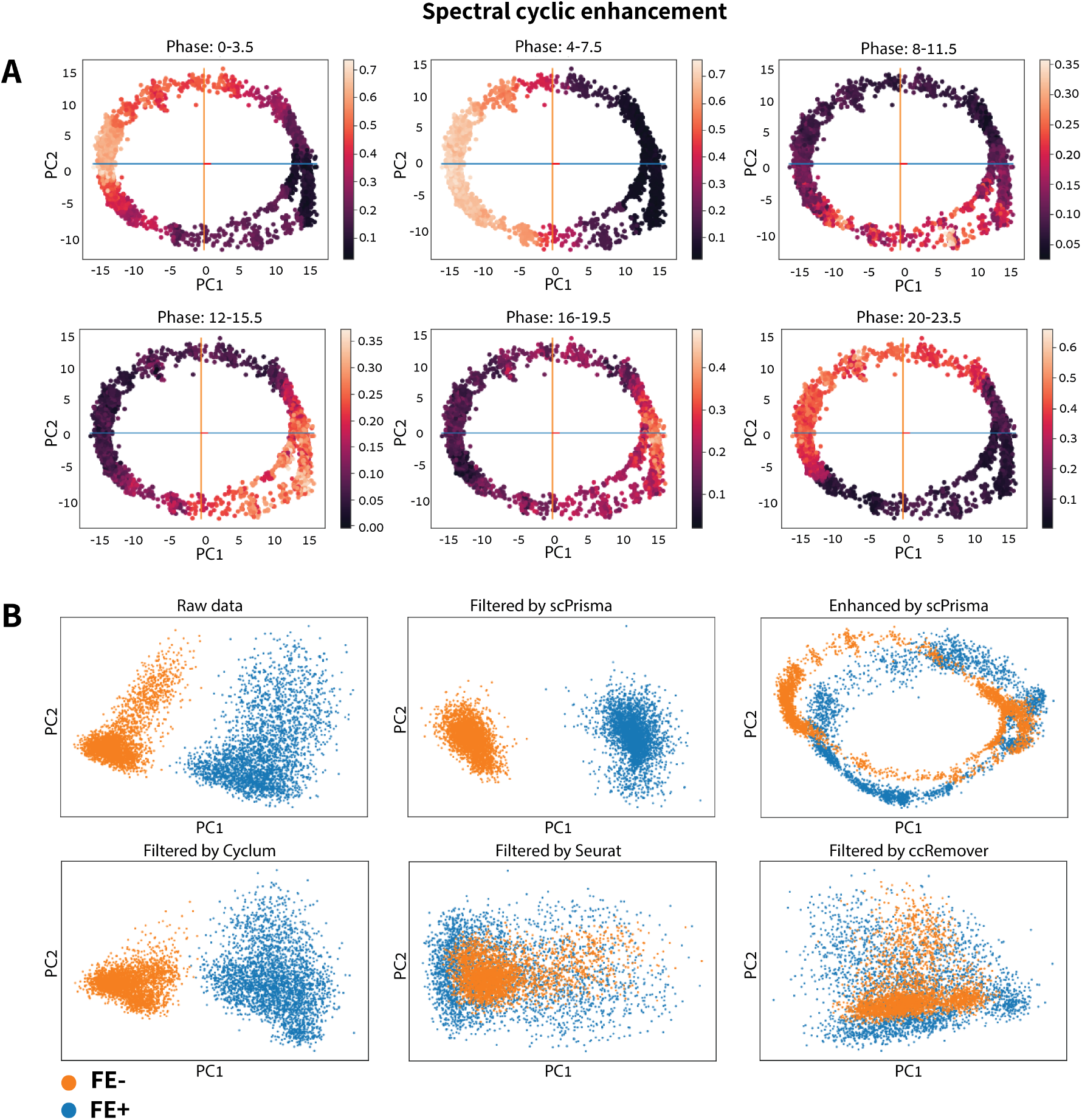
scPrisma detects and filters the diurnal cycle in Chlamydomonas. (A) PCA representation of cells in spectrally enhanced cyclic signal of the iron replete (Fe+) experiment [27]. Each plot represents 1/6 of the 24 hours cycle. The cells are colored according to the normalized sum of the marker genes associated with each phase [44]. (B) PCA representation of cells in iron replete (Fe+, blue) and iron deficient (Fe-, orange) conditions of raw gene expression data, data following scPrisma’s cyclic filtering (Silhouette score before/after filtering = 0.088/0.136, Davies-Bouldin score before/after filtering = 3.118/2.422, Calinski and Harabasz score before/after filtering = 600.802/1000.44) and enhancement, as well as data following filtering by Cyclum (Silhouette score = 0.052, Davies-Bouldin score = 4.183, Calinski and Harabasz score = 337.629), Seurat (Silhouette score = 9.868e-04, Davies-Bouldin score = 136860121.54, Calinski and Harabasz score = 1.29e-10), and ccRemover (Silhouette score = 9.815e-06, Davies-Bouldin score = 6759170.71, Calinski and Harabasz score = 6.25e-13).

### 2.6 scPrisma extracts cell-type specific circadian rhythm signals in the suprachiasmatic nucleus

Finally, we focus on scRNA-seq data collected for mice suprachiasmatic nucleus (SCN), the mammalian brain’s circadian pacemaker [26]. In this experiment, cells were sampled at 12 different timepoints along two days. Again, we leveraged the cyclic nature of the circadian rhythm, and explicitly used the prior knowledge regarding the experimental sampling times (instead of running the reconstruction algorithm). We first clustered the cells using Louvain algorithm, and mapped individual clusters to cell types using established marker genes[26] (Supplementary A.6, Supplementary Figure 5). When analyzing each cell type separately, we were able to reveal a cyclic signal associated with the circadian rhythm for 5/8 of cell types using scPrisma’s cyclic enhancement algorithm (Figure 6, Supplementary Figure 6, Methods 4.4). The three cell types that did not expose a clear cyclic signal following cyclic enhancement (NG2, microglia, and tancytes) are indeed those which exhibit the lowest fraction of rhytmic genes expression out of total expressed genes [26] (Supplementary Figure 6). Moreover, we used the Calinski and Harabasz score[4] to measure the separation of cells that were sampled at different time points, before and after cyclic filtering/enhancement. We find that overall, as expected, separation increases substantially following cyclic enhancement and decreases following cyclic filtering, which as above, is least substantial for the three cell types exhibiting the lowest fraction of rhythmic genes (Figure 6D). It can be observed (Figure 6A) that cellular density varies across the different time points, which seems to reflect underlying biological variability as time points that were previously identified as peaks of the rhytmic process[26] exhibit higher cellular concentration. For example, for ependymal cells, the timepoints CT06, CT10 and CT14 exhibit high cellular concentrations (Figure 6A), while being reported as the peaks of the rhythmic process using gene ontology[26], consistent with rhythmic genes mainly expressed at these timepoints, such as *Tef* which is known to play a role in circadian regulation[48], and is mainly expressed around CT10 and CT14 (Figure 6B).

**Figure 6:**
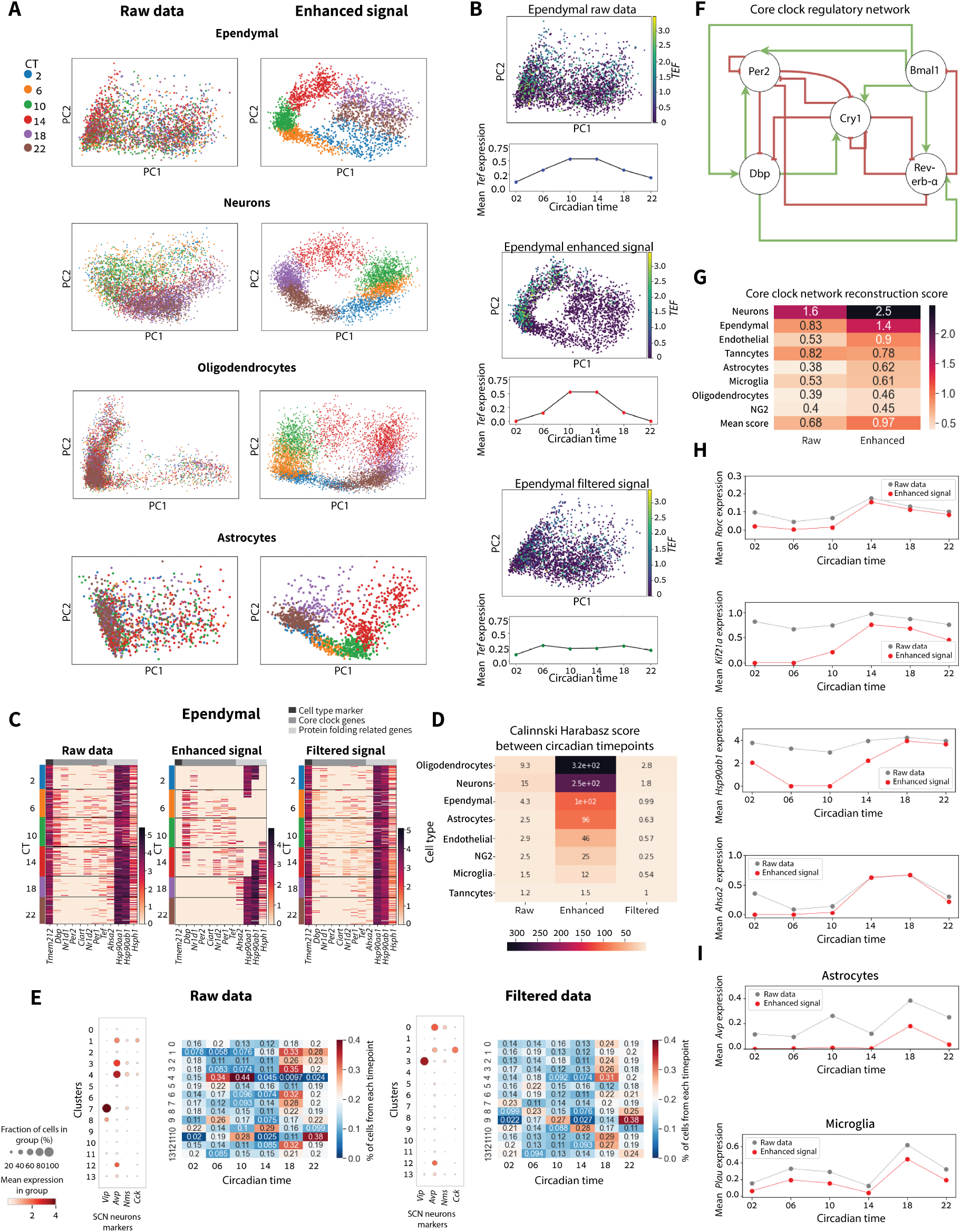
scPrisma extracts cell-type specific circadian rhythm signals in the suprachiasmatic nucleus. (A) PCA representation of cells in raw SCN gene expression data [26] and data following cyclic enhancement by scPrisma, colored according to circadian timepoints (CT), for several cell types: ependymal, neurons, oligodendrocytes and astrocytes. (B) PCA representation of ependymal cells colored by *Tef* expression and mean *Tef* expression as a function of circadian time, for raw data (top), data following scPrisma’s cyclic enhancement (middle) and filtering (bottom). (C) Heatmaps of ependymal expression of (from right to left) cell type marker genes, rhytmic genes and protein folding genes, for raw data (left), and data following scPrisma’s cyclic enhancement (middle) and filtering (right). (D) Calinski Harabasz score for cells sampled at different circadian timepoints for raw data, and data following scPrisma’s cyclic enhancement and filtering. Cell types with low fraction of circadian genes[26] exhibit lower scores. (E) Pre- and post-filtering gene expression dotplots of the marker genes of SCN neurons (prior to filtering, clusters 1, 3 and 4 contain mixtures of two different subtypes), as well as pre- and post-filtering heatmaps showing the contribution of cells sampled at different circadian time points to each neuronal cluster. (F) Regulatory network structure of core clock genes[34]. (G) Pre- and post-enhancement sum of scores of all regulatory interactions of the core clock network shown in (F), scored by GRNBoost2, for each SCN cell type. (H) Pre- and post-enhancement mean *Rorc, Ahsa2*, and *Hsp90ab1* expression as a function of circadian time point. Following cyclic enhancement, regulatory interactions between the transcription factor *Rorc* and *Ahsa2, Hsp90ab1* were uncovered. (I) Mean *Avp* and *PLAU* expression as a function of circadian time point. Following cyclic enhancement, cell-cell interactions between astrocytes and microglia cell types, mediated by *Avp* and *PLAU*, were revealed.

Focusing on gene expression, we find that spectral cyclic enhancement indeed diminishes the expression of cell type marker genes and retains the expression of rhythmic genes (core clock genes and protein folding genes, as characterized in [26]) (Figure 6C, Supplementary Figure 6). Conversely, following cyclic filtering, cell type marker gene expression is retained while the resulting expression of rhythmic and protein folding-related genes become temporally uniformly distributed (Figure 6C, Supplementary Figure 6).

Next, we will exemplify how spectral enhancement by scPrisma allows us to achieve better cell type classification, infer gene regulatory interactions related to the circadian rhythm, and reveal underlying cell-cell interactions. When aiming to characterize cells by their type, additional biological signals which are encoded in the cellular gene expression can interfere with that task, as similarity between cells can arise due to multiple different factors. For example, direct clustering of cells according to their single-cell gene expression profiles may capture similarity according to circadian rhythm phase, and not necessarily their type, which can substantially hinder our ability to distinguish between different cell types. This is in fact the case for the SCN neurons analyzed above. When clustering the neurons, the outcome is 13 distinct clusters, 4 of which can be identified using either established marker genes or subtype classification in a previous study[26] as SCN neuronal subtypes (Supplementary A.6, Figure 6E). However, two of the clusters are each correlated with two previously-established subtype expression profiles (Supplementary A.6, Figure 6E). We find that the circadian rhythm signal interferes with the proper classification of subtypes in this case, supported by the observation that in the first of the two mixed clusters the majority of cells (79%) were sampled between CT =14/18/22 while in the second mixed cluster the majority of cells (93%) were sampled in CT = 02/06/10, which suggests that the clustering of cells in this subpopulation may be dominated by their distinct temporal signature and not necessarily by their type (Figure 6F). We were able to overcome the cell type misclassification by spectrally filtering out the circadian rhythm using scPrisma, after which, the clustering algorithm yielded a unique cluster for each neuronal subtype (Figure 6E). Moreover, as expected, the distribution over circadian timepoints within each cluster flattened following cyclic filtering (Figure 6E; mean circular variance increased from 0.781 to 0.863 following filtering).

Mixed biological signals in single-cell data can also interfere with the inference of gene regulatory networks. Therefore, we used scPrisma to highlight a set of regulatory interactions related to the circadian rhythm that were difficult to identify in the original data. Specifically, we expect that regulatory interactions that can be revealed following cyclic enhancement of the single-cell data (and are hidden in the original data) would be enriched with interactions associated with the cyclic circadian process. Indeed, regulatory interactions between core clock genes, as inferred using both a gene regulatory network inference algorithm (GRNBoost2[29]) and a simple gene-gene covariance analysis (Supplementary A.6), are more highly correlated to the established core clock interaction network[34] (Figure 6F) in the cyclically enhanced single-cell data, relative to the raw data, for 7 of 8 of the SCN cell types (Figure 6G, Supplementary A.6). Going beyond the known core clock interaction network, we searched for inferred interactions (based on GRNBoost2) which are substantially enhanced following scPrisma cyclic analysis (Methods), where the regulator is a core clock transcription factor (*Nrld1, Nrld2, Rora, Rorb, Rorc, Dbp, Tef*) [19]) (a full list of inferred interactions is available in supplementary Table 1). For example, focusing on genes inferred to be highly regulated by *Rorc* in ependymal cells following spectral enhancement, we find that the genes that received the highest score are: *Ahsa2* (0 to 6.341), *Kif21a* (0 to 6.231), *Hsp90ab1* (0.070 to 5.731) and *Mt1* (0 to 5.002), and the peaks of these genes along the circadian rhythm overlap with the peak of *Rorc*, following spectral enhancement (Figure 6H and Supplementary Figure 5E). These results are consistent with previous results showing the existence of a regulatory interaction between *Rorc* and *Hsp90ab1* which is dependent on the time of day [23].

Finally, we use scPrisma to infer hidden cell-cell interactions related to the circadian rhythm. We compared cell-cell communication patterns, based on inferred ligand-receptor interactions using Cell-PhoneDB database and method [8], between different cell types at corresponding time points. Similarly to the regulatory network inference described above, we searched for interactions that were substantially enhanced following scPrisma’s cyclic analysis (Methods 4.7) (The full list of predicted cell-cell interactions can be found in Supplementary Table 2). Such enhanced interactions are expected to be enriched for those associated with the circadian rhythm, or rhythmic processes more generally. We specifically searched for interactions whose score increased at least two-fold following scPrisma’s enhancement (Supplementary A.6). An example of such enhanced inferred interaction is between *Avp* in astrocytes and *PLAU* in microglia, both known to be associated with the circadian rhythm [32, 33], whose mean score in CT18 was increased following enhancement from 0.501 to 1.011 (p-value=0.003 before enhancement, p-value < 0.001 after enhancement) (Methods, Figure 6I). More generally, we searched for interactions associated with rhythmic processes using the corresponding gene ontology categorization (rhythmic process, GO:0048511)[16], and termed cell-cell interactions mediated by a receptor (ligand) which is associated with a rhythmic process as receptor (ligand) -mediated rhythmic interactions. The percentage of inferred receptor-mediated rhythmic interactions which were substantially enhanced (at least 2-fold, as above) following cyclic enhancement increased to 16.32%, relative to 11.61% before enhancement. Similarly, the percentage of ligand-mediated rhythmic interactions increased to 14.28% relative to 9.68% before enhancement. In summary, the results for predicted gene regulatory interactions and cell-cell interactions following cyclic enhancement by scPrisma we describe above (Supplementary Tables 1,2) are expected to reveal and strengthen previously overshadowed interactions related to cyclic processes such as the circadian rhythm, which is supported by the results over the subset of known core clock regulatory interactions, as well as enrichment of rhythmic genes among ligands and receptors of predicted cell-cell interactions.

## 3 Discussion

In this study, we developed scPrisma, a spectral analysis workflow for pseudotime reconstruction, informative genes inference, signal filtering and enhancement. scPrisma presents three major contributions; First, it embodies a full workflow for analyzing underlying periodic and linear signals based on a topological approach that can be performed either de novo or enhanced using prior knowledge integration (e.g. low-resolution pseudotime or marker gene information). This flexibility allows scPrisma to uncover cyclic signals of varyingstrengths. Second, scPrisma enables both signal enhancement and filtering without embedding to lower dimensions, which makes scPrisma useful as a prior step for multiple types of existing downstream analyses. This can accelerate biological discovery, as we exemplify above for SCN neurons, by revealing gene regulation patterns and cell-cell interactions that are associated with a specific biological process such as the circadian rhythm, or clustering cell groups according to a specific characteristic, such as cell type, while filtering similarity due to time of day. Third, the enhancement algorithm does not overfit to a circular topology and does not force a circular topology when applied to expression data encoding no or negligible rhythmic signals. scPrisma avoids overfitting by: applying the genes inference task before the enhancement (and thus retaining only genes related to the desired signal), controlling the level of filtering by regularization, and by restricting the range of entries in the filtering matrix.

A computational challenge arises due to the non-convexity of the reconstruction and enhancement tasks. This optimization challenge is relieved due to the cyclic topology prior, as the theoretical analysis of the eigenvectors does not depend on the specific values of the matrix, but only on its circulant property. In addition, the multiple solutions for the cyclic reconstruction task (every circular, clockwise or counterclockwise, shift of a solution is also a solution), ease the convergence to a feasible solution.

Future work can extend scPrisma to more topologies, beyond cyclic and linear, by tailoring a covariance matrix for any desired topology, and using scPrisma’s existing workflow for analysing the desired underlying signals. We anticipate that scPrisma will accelerate single-cell based research by enhancing target signals of interest and enabling their identification and analysis, and providing a general workflow for single-cell signal disentanglement in diverse biological contexts.

## 4 Methods

### 4.1 Spectral analysis of cyclic signals

For theoretical analysis, we constructed three simple models for the cyclic signals. In the first model, we receive as input the number of cellular variations (‘*n*’), the numbers of genes (‘*p*’), and the number of changes between neighboring cells (‘*k*’). We start with a ‘root’ cell whose expression profile is a binary vector with p entries ({1, 0}^*p*^). Each gene is approximated to be either expressed (ON,’1’) or not expressed (OFF,’0’). Then, the next cell in the cycle is generated by duplicating the existing cell, choosing uniformly ‘*k*’ genes and switching their state. This process is repeated ‘*n*’ times. Then, within the last generated cell, ‘*k*’ genes whose state differs from the ‘root’ cell are chosen at random, their state is switched and the cell is duplicated. This process is then continued until the gene expression of the newest cell is identical to the ‘root’ cell. Illustration can be seen in (Supplementary Figure 2).

For analyzing the covariance matrix of the model, we will use similar Markovian assumption to the assumption that was used in Nitzan & Brenner, 2021 [30] and Qin & Colwell [36]; the covariance between the expression profiles of two cells, separated by *m* state change, where *m* is the minimum distance between the cells clockwise and counterclockwise (cyclostationary assumption), is given by *α*(*m*) = *E*[*X*(*m*)*X*(0)] = exp(−2*mk/p*). According to this assumption, the expected covariance matrix of the gene expression matrix is circulant:

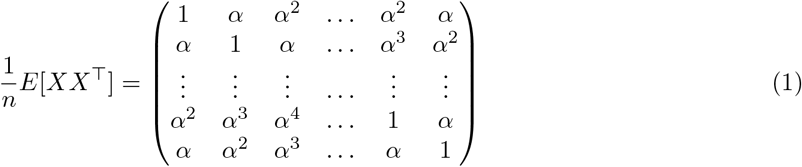

The (*k,j*) entry of a general circulant matrix *C* is given by *C*_*k,j*_ = *c*_(*j*−*k*)%*n*_ [13]. The spectrum of a circulant matrix has analytical closed formula [37]. Specifically, the eigenvalues are the discrete Fourier transform of the first row, and the eigenvectors are the normalized Fourier modes. Since a covariance matrix is symmetric and positive semi definite, all its eigenvalues are real. Therefore the eigenvalues are the discrete cosine transform of the first row [6]:

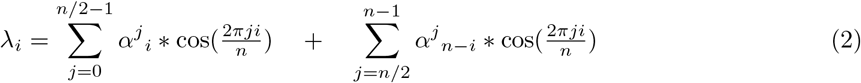

The sth entry of the *i* th eigenvector corresponding to the ith eigenvalue is [6]:

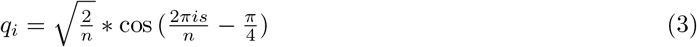

For testing our approach we defined two additional models are described in Supplementary methods A.1, we used this models in the Simulated data section (subsection 2.2).

### 4.2 Spectral analysis of linear signals

Similarly to the analysis of cyclic signals, we first construct a simple model for the linear signals. We follow a similar linear model to the one that was presented by Nitzan & Brenner [30]. The model receives the same input as the cyclic model, and each cell is represented with a binary vector of ({1, 0}^*p*^). We start from a ‘root’ cell, and then over n iterations, a new cell is created in the linear chain by changing the state of ‘*k*’ randomly chosen genes relative to the previous cell in the chain. As in the cyclic model, we assume that the covariance between the gene expression profiles of two cells, separated by ‘*m*’ state changes, is given by *α*(*m*) = *E*[*X*(*m*)*X*(0)] = exp (–2*m/p*). Thus, the expected covariance matrix of the gene expression matrix is:

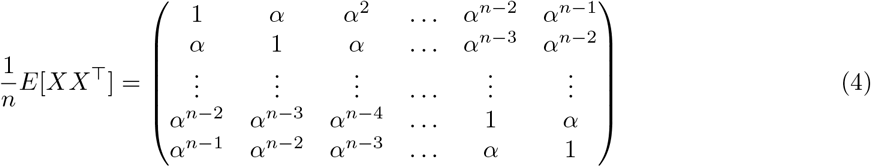

This matrix is a special case of a Toeplitz matrix, and is particularly known as “Kac-Murdock-Szego” matrix [14]. The eigenvalues of such a Kac-Murdock-Szego matrix can be approximated as [14]:

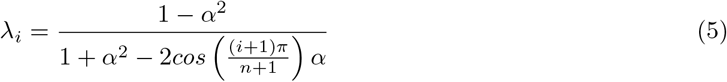

The corresponding eigenvectors can either be estimated analytically[45] or calculated by the numerical decomposition of the theoretical matrix.

### 4.3 Pre-processing

We used standard pre-processing pipeline. First removing genes that are not expressed in any of the cells in our data, applying per-cell normalization by dividing each count by the total counts of the particular cell, applying log transformation, and retaining only highly variable genes.

For the reconstruction algorithm, scaling the *L*_2_ of each cell to 1, should be applied, to ensure that the circulant matrix has constant diagonal. For the gene inference algorithm, scaling the *L*_2_ of each gene to 1 should be applied, as the score of each gene is relative to the rest of the genes. For estimating *α* (that represents the correlation between neighbors according to the target topology), we search for the *α* value that best matches the spectrum of the given gene expression matrix. The results were improved by applying the algorithms after removing the theoretical covariance vector associated with the largest eigenvalue. For the cyclic case, the values of this eigenvector are constant.

### 4.4 Algorithm

1. Choose the desired topology (periodic/linear). Calculate a theoretical covariance eigenvectors and eigenvalues.
2. Pre-process the data.
3. Reconstruct the signal by reordering the gene expression rows by solving Problem 2 (below), or by using prior knowledge.

Option 1 - signal enhancement:

a. Infer informative genes by solving Problem 3 (below), or by using prior knowledge, and remove the rest of the genes.
b. Enhance the desired signal by solving Problem 4 (below).

Option 2 - signal filtering:

a. Filter out the desired signal by solving Problem 5.

### 4.5 Signal reconstruction

With a closed formula for the spectrum (2,3), we can estimate the pseudotime of the underlying cyclic trajectory. This can be done by estimating the rows reordering of the gene expression matrix that maximizes the projection over the theoretical spectrum. This problem can we formulated as a matrix permutation problem.

#### Problem 1.

*Matrix permutation problem for estimating pseudotime that maximizes the projection over the theoretical spectrum*

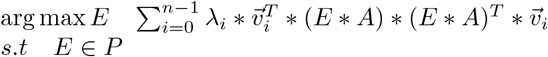

Where *A* is the original gene expression matrix, 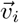 and *λ_i_* are the theoretical eigenvectors and corresponding eigenvalues, respectively, and *P* is the set of permutation matrices. Under the assumption that A has a permutation 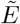 such that the spectrum of 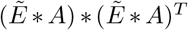 matches that of the theoretical spectrum, the optimal solution is 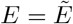 (see Supplementary proof 1). Consider an identical formulation of the function we wish to maximize: 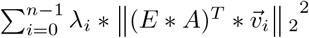. This formula aims to maximize the product of each gene sorted by the permutation matrix and each theoretical eigenvector multiplied by its eigenvalue. These theoretical eigenvectors, as they are the eigenvectors of the theoretical covariance matrix, represent the variance along the theoretical topology. As a result, the permutation which maximizes this objective maximizes the variance along the theoretical topology.

Permutation problems are known to be NP-Hard[10]. Convex relaxation was used in previous studies, and instead of searching a permutation matrix, a doubly stochastic matrix (the Birkhoff polytope) was searched[10, 46].

#### Problem 2.

*Convex relaxation for Problem 1*

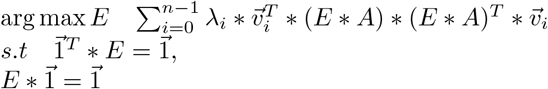

The objective function is quadratic and convex (see Supplementary proof 2). Despite the fact that maximizing it is not convex optimization, previous studies have shown that such problems can be efficiently resolved with stochastic gradient descent [42] [49]. For projection into “Birkhoff polytope”, we used Bregmanian Bi-Stochastication algorithm [46]. Finally, for rounding the doubly stochastic matrix to permutation matrix we used a simple greedy algorithm. Specifically, the algorithm iterates over all rows, for each row rounds the maximum entry in each column that doesn’t have ‘1’ value yet to ‘1’, and rounds to ‘0’ the rest of the entries. The output of this algorithm is the permuted gene expression matrix: *A_ordered_* = *E* * *A*.

### 4.6 Genes inference

Once the reconstructed signal is obtained, either by solving Problem 2 or by prior knowledge, identification of informative genes that are related to the desired signal, is possible. This can be achieved by filtering genes that do not show spatial behavior associated with the reordering, or genes that do not maximize the projection over the theoretical spectrum. Because of convexity considerations, it would be easier to infer genes that are not related to the desired signal, and then flip the results. Inference of the genes that are not related to the desired signal can be done by solving the following optimization problem:

#### Problem 3.

*Genes inference*

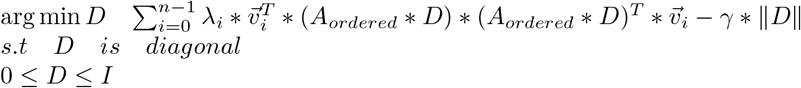

Since genes are represented by columns of *A*, each entry on the diagonal of *D* represents the influence of the respective gene on the spectrum. The number of filtered genes can be controlled by adding regularization. We can either increase the regularization coefficient (γ) to filter fewer genes or decrease it to filter more genes. The output of this algorithm is the gene expression matrix, after nullifying the genes that are not informative relative to the signal: *D*_1_ = *I* – *D, A_genes inferred_* = *A_ordered_* * *D*_1_.

### 4.7 Filtering and enhancement

After inferring the set of informative genes (the genes related to the reconstructed signal), the next step is to remove any information that is not related to the signal, from the expression of those genes. This can be achieved by removing the diagonal constraint from Problem 3, and replacing the matrix product by Hadamard product (element-wise product). For every entry in the expression matrix (*A_i,j_*), this formulation matches an optimization variable (*F_i,j_*). Therefore, the enhanced gene expression matrix contains only expression profiles that maximize the projection over the theoretic spectrum.

#### Problem 4.

*Signal enhancement*

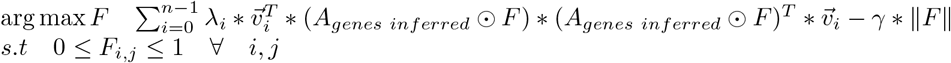

Since this problem is not a convex optimization problem, it can be solved by using stochastic gradient ascent (adding noise at each iteration [5]). The output of this algorithm is the gene expression matrix, after eliminating information that is unrelated to the signal of interest: *A_enhanced_* = *A_genes inferred_* ⊙ *F*.

Another option is similar to that described for Problem 3, which is to transform this problem to a minimization problem for filtering the reconstructed signal, thus turning it into a convex optimization problem (see Supplementary proof 2). Formulating this problem as a minimization problem, enables to eliminate the variance along the theoretical topology.

#### Problem 5.

*Signal filtering*

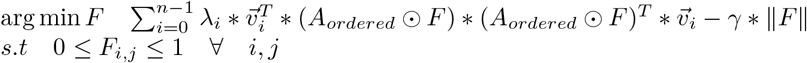

The output of this algorithm is the gene expression matrix, after eliminating the information that is related to the signal of interest: *A_filtered_* = *A_ordered_* ⊙ *F*.

## 5 Code availability

The code is publicly available at: https://github.com/nitzanlab/scPrisma/

## 6 Data availability

The scRNA-seq datasets used for this study were acquired from the Gene Expression Omnibus (GEO) database with the following accession numbers: HeLaS3 ([GSM4224315](https://www.ncbi.nlm.nih.gov/geo/query/acc.cgi?acc=GSE142277)), liver (GSE145197), Chlamydomonas (GSE157580) and SCN (GSE117295).

## A Supplementary Notes

### A.1 Cyclic models

In the first model - the progression model (which is described in the main text), we start with a ‘root’ cell. Next, we duplicate the existing cell, select uniformly ‘*k*’ genes, and switch their states. This process is repeated ‘*n*’ times. Then, within the last generated cell, ‘*k*’ genes whose state differs from the ‘root’ cell are randomly chosen, their state is switched and the cell is duplicated. The process is repeated until the gene expression of the newest cell is identical to the original root cell.

In the second model - the spatial model, we receive as input the number of cells (‘*n*’), the numbers of genes (‘*p*’) and a window size (‘*w*’, where 0 < *w* < 1). The simulation starts by arranging the ‘*n*’ cells on a circle and initializing the gene expression of each one of them as ‘OFF’ for all genes. Then, for each gene we choose uniformly the center cell and switch the gene to ‘ON’ for the cells inside the window of size *w * n* around the center cell. Illustration can be seen in (Figure 1B). The third model is similar to the second model, but instead of step functions of gene expression around each center cell, expression is estimated by a Gaussian distribution centered at the center cell.

### A.2 Simulated data

For the reconstruction evaluation, we simulated a cyclic signal according to model B (Supplementary A.1). The signal consists 100 cells and 500 genes with w=0.3. We did not shuffle the rows, so the ground truth order is the identity permutation (or any other shift or shift of the reverse order). For 300 iterations we added Gaussian noise with varying variance, and used the reconstruction algorithm. Next, the permutation matrix was transcribed to permutation array. And finally, the maximum Spearman rank correlation was taken between the output permutation array and every possible shift and inverse shift of the identity permutation.

For the genes inference evaluation, cyclic signal of 256 cells and 250 genes was simulated according to model B, And a lineage signal of 256 cells and 250 genes was simulated according to the model described by Nitzan & Brenner [30]. Again, for 300 iterations we added Gaussian noise with varying variance, and used the genes inference task. As ground truth vector cyclic genes were labeled as ‘1’ and the lineage genes were labeled as ‘0’. The AUC-ROC of the diagonal of ′*D*′ matrix and the ground truth vector was calculated.

### A.3 HeLa cells data

We used similar pre-processing as in [50], including normalization per cell, log transformation and selecting only the highly variable genes (7000). Additionally, similarly to [41], we filtered out cells with low number of counts, until the mean counts reached 4,500 UMI counts per cell. The polar plot (Figure 3B) consists of 5 different phases (G1.S, S, G2, G2.M, M.G1), where for each phase we used the list of genes found to be associated with it [41]. For each phase, we created a vector that represents the probability distribution of each cell to be in each cell cycle phase. For calculating those vectors, we summed all the phase-related genes and then normalized the sum of each vector to 1. For evaluating the performance of the algorithms, we measured two terms for each vector: circular mean 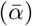 and circular variance[17]. In angular statistics, circular mean is a measure that reflects the mean of a set of angles or similar cyclic quantities. Given a set of observation of angles: *α*_1_,…*α_n_*, the circular mean is 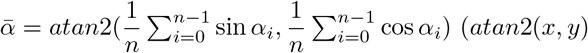 returns a value *θ* between – *π* ≤ *θ* ≤ *π*, where for some *r* > 0: *x* = *r* cos *θ, y* = *r* sin *θ*). In our case, given an ordering of the cells in a cycle (according to the reconstruction algorithm), we have *n* (the number of cells) samples that are distributed uniformly along the cycle, but the probability (the weight) of each one of them is different, and the total weights are summed to 1. So the circular mean would be: 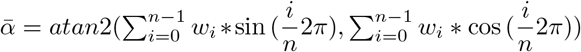, where *w_i_* is the i’th entry in the expression vector (related to the i’th cell). Circular variance measures the spread of dihedral angles. The circular variance ranges from 0 to 1, where lower values reflect clustering of the samples around the circular mean The circular variance is defined as 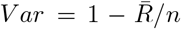, where 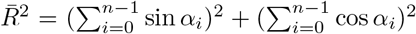. In our case, since the distribution is not uniform we will calculate: 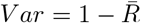, where 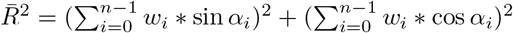.

For evaluating the informative genes inference task, we used the same list of genes. We chose random subsets of 100 genes from the list, and 100 genes that do not appear on the list (50 iterations). The genes inference was conducted over the 200 genes that were chosen.

### A.4 Liver data

We used standard pre-processing (normalization per cell, log transformation and selecting the highly variable genes). For labeling the cells according to their position along the lobule axis, we used the algorithm in [7]. The output of this algorithm is a probabilistic embedding, where each cell was eventually labeled by the layer associated with the highest probability.

### A.5 Chlamydomonas data

To verify that our reconstruction corresponds to the diurnal cycle, we used a list of marker genes that was obtained from bulk RNA-seq[44], where the peak time of genes that are influenced by the diurnal cycle are annotated. We divided the 24-hours diurnal cycle to 6 equally-spaced segments (0-3.5,4-7.5,8-11.5,12-15.5,16-19.5,20-23.5). Then, similarly to the HeLa cells analysis (Supplementary A.3), the genes were summed and normalized by dividing all values by the maximum value.

### A.6 SCN data

We clustered the cells according to cell types by using Louavin algorithm, where the mapping between clusters to cell types was done using established cell type marker genes[26]. We used *AGT* for astrocytes, *ITM2A* for endothelial, *TMEM212* for ependymal, *HEXB* for microglia, *CELF4* for neurons, *PDGFRA* for NG2, *PLP1* for oligodendrocytes, and *col23a1* for Tanyceytes. The expression of each marker gene in each cluster is shown in Supplementary Figure 5.

We recovered the neurons subtypes by clustering using the Leiden algorithm. According to [26], there are four main SCN neuron subtypes: ‘N0’ (marker genes: *Avp/Nms*), ‘N2’ (marker genes: *Avp, Cck*; can be divided into two subgroups), ‘N6’ (marker genes: *Vip, Nms, C1QL3*), and ‘N9’ (marker genes: *Vip, Grp*). Focusing on N0 and N2, When running a naive clustering algorithm (Leiden) over the cells that were classified as neurons, we get 13 clusters. The marker genes of N0 and N2 are expressed in 4 of these clusters: 1,3,4 and 12 (where cluster 12 contains only 293/14766 cells). We conclude that clusters 1,3, and 4 contain a mixture of N0 and N2 cells, since all three marker genes (*Avp/Nms/Cck*) are expressed in all three clusters. Moreover, the majority of cells in clusters 1 and 3 (79% and 66%, respectively) are sampled in the timepoints CT =14/18/22, while the majority of cells in cluster 4 (93%) are sampled in the timepoints CT = 02/06/10. Only after cyclic filtering, can cluster 0 be uniquely classified as N0 (according to *Nms* expression), while cluster 2 can be classified as N2 (according to *Cck* expression) (Figure 6E).

We evaluated the prediction of regulatory interactions associated with the circadian rhythm, using the GRNBoost2 algorithm, relative to a baseline circadian network [29]. For each cell type separately, we summed the scores of all regulatory interactions in the circadian network. In addition, we calculated the correlation between the expression levels of each pair of interacting genes according to the circadian network. We scaled the expression of the circadian network genes (mean=0 and std=1) and calculated the covariance between interacting genes.

### A.7 Comparisons

For comparing the filtering and the enhancement of the simulated data vs Cyclum[24], we used the defualt parameters of Cyclum (with the same pre-processing as scPrisma), except for the dimensionality of the embedding layer. For finding the optimal embedding layer, we ran it with eleven different values for dimensions {1, 2,…11} as suggested in the original paper, and we chose the best result.

For comparing the filtering of the diurnal cycle, we used Cyclum as described for the simulated data. For comparing with Seurat [3] and ccRemover [1]), we used the marker genes list of the diurnal cycle that was provided by[44]. For ccRemover we used the whole list. For Seurat we sorted the genes according to their peak phase, randomly divided the genes list to two ranked subsets randomly (over 50 iterations) and chose the best result.

### A.8 Regularization parameter

In our method, the regularization parameter (*γ*) is the only significant free parameter, and incorrect selection of this parameter can result in non-optimal filtering. We recommend using *γ* = 0 as the starting point for all the algorithms. If the enhancement algorithm does not filter out enough information, it is recommended to increase the regularization parameter incrementally by 1/*n*, where *n* is the number of cells. For the filtering algorithm, if the algorithm filters out too much information, it is recommended to increase the regularization parameter incrementally by 1/*n*. For the gene inference task, we recommend increasing the regularization parameter incrementally by 0.1 until a suitable number of genes are retained.

### A.9 Integrating prior knowledge for the reconstruction problem

Since the reconstruction task is challenging, integrating any available prior knowledge can be helpful, such as restricting the optimization parameter (E) to a convex subset of the doubly stochastic matrices set. For example, we simulated a cyclic signal with varying amount of noise, and applied the reconstruction algorithm both without prior knowledge (de novo) and with prior knowledge related to low-resolution pseudotime ordering (specifically, division of the cells to three consecutive temporal bins).We find that integration of prior knowledge in this case renders the algorithm robust to high noise levels (*SNR* < 0.1), as opposed to the de novo version (Supplementary Fig 2E).

Another possible type of prior knowledge that can be integrated is related to gene selection, where we can apply the reconstruction algorithm only over a subset of genes that are known to be related to the desired signal. For example, in many scRNA-seq datasets, cell type is a major source of variation reflected in the covariance matrix, and therefore, better results are generally expected if the reconstruction algorithm of the cell cycle is applied after selecting for genes that are known to be related to this process.

### A.10 Proof of correctness

#### Theorem 1.

*Let A * A^T^ (where A ∈ R^n×m^) be a circulant matrix with eigenvalues λ*_1_, *λ*_2_…*λ_n_, and eigenvectors* 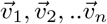, *and assuming that we have Ã which is a rows permuted version of A, then E * Ã* * (*E * A*)*^T^ is circulant, where E is the solution of Problem 1*.

*Proof*. It is sufficient to show that given two permutation matrices *E* and 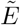, if *E * Ã* * (*E * Ã*)^*T*^ is circulant with the eigenvalues *λ*_1_, *λ*_2_…*λ_n_*, and eigenvectors 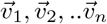, then:

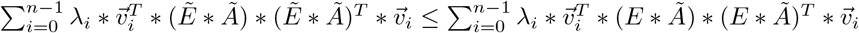

Assuming that the eigenvectors of 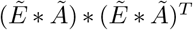 are 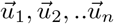, and since eigenvalues are insensitive to permutations, the eigenvalues are *λ*_1_, *λ*_2_…*λ_n_*.

Left side:

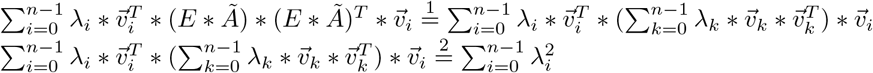

Right side:

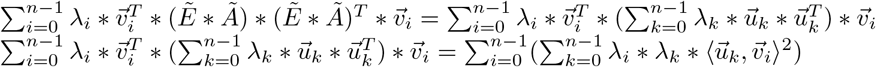

Therefore we need to show that: 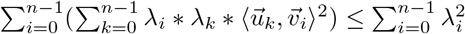.

Ky Fan [9] (LEMMA 1A) proved a similar lemma:

Let *a*_1_ ≥ *a*_2_ ≥ ….*a_n_* ≥ 0, *b*_1_ ≥ *b*_2_ ≥ ….*b_n_* ≥ 0. If *p_ij_* are *n*^2^ non-negative numbers such that:

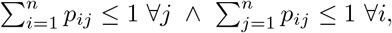

then:

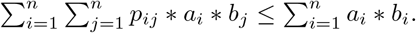

In our case, we can use the lemma since from Bessel’s inequality: 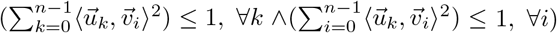, and all the eigenvalues are non-negative since the covariance matrix is a PSD matrix. Therefore, we can use the lemma with: *a_i_* = *λ_i_, b_j_* = *λ_j_*∀*i, j*.

### A.11 Proof of convexity

To prove the convexity of the objectives of the optimization problems (Methods 4.5, 4.6, 4.7), we will use the following theorem:

#### Theorem 2.

*f*: *R* → *R^n^ is convex if and only if g*: *R* → *R is convex. where g*(*t*) = *f*(*x+tv*), **dom** *g* = {*t*|*x* + *tv* ∈ **dom** *f*}*[2]*

**Claim 2.1.** *The objective function of the convex relaxation of the reconstruction problem (Methods 4.5) is convex*.

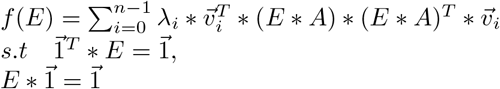

*Proof*. The set iself is convex (linear equalities). Let us define *g*(*t*) = *f*(*E* + *tU*), where *E* + *tU* ∈ **dom***f*, and calculate the first and second derivatives:

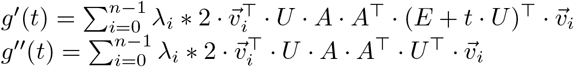

Since *U* · *A* · *A*^T^ · *U*^T^ is positive semi-definite, the second derivative of *g*(*t*) is positive ∀*t*. Therefore *g* is convex, and according to theorem 2, f is convex as well.

**Claim 2.2.** *The objective function of the genes inference problem (Methods 4.6) is convex (and since it is a minimization problem, it is a convex optimization problem)*.

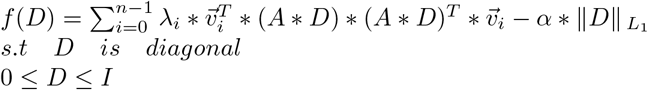

*Proof*. The set itself is convex (linear inequality). Similarly to claim 2.1, we will define *g*(*t*) = *f*(*E* + *tU*), where *E* + *tU* ∈ **dom***f*, and calculate the first and second derivatives:

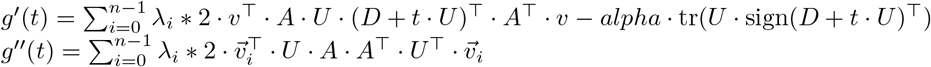

Since *U* · *A* · *A*^T^ · *U*^T^ is positive semi-definite, the second derivative of *g*(*t*) is positive ∀*t*. Therefore, *g* is convex, and according to theorem 2, f is convex as well.

**Claim 2.3.** *The objective function of the enhancement and filtering problems (Methods 4.7) is convex (and since the filtering problem is a minimization problem, it is a convex optimization problem)*.

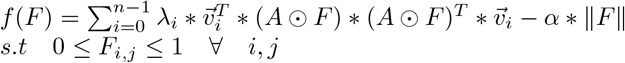

*Proof*. Under the assumption of *A* being a gene expression matrix with positive entries, we will transform the objective and the set to be:

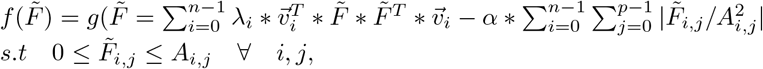

where 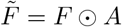. After the rewriting, we will define 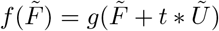:

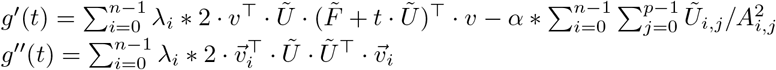

Similarly to the previous claims, since *U* · *A* · *A*^T^ · *U*^T^ is positive semi-definite, the second derivative of *g*(*t*) is positive ∀*t*. Therefore *g* is convex, and according to theorem 2, f is convex as well.

## B Supplementary figures

**Supplementary Figure 1:**
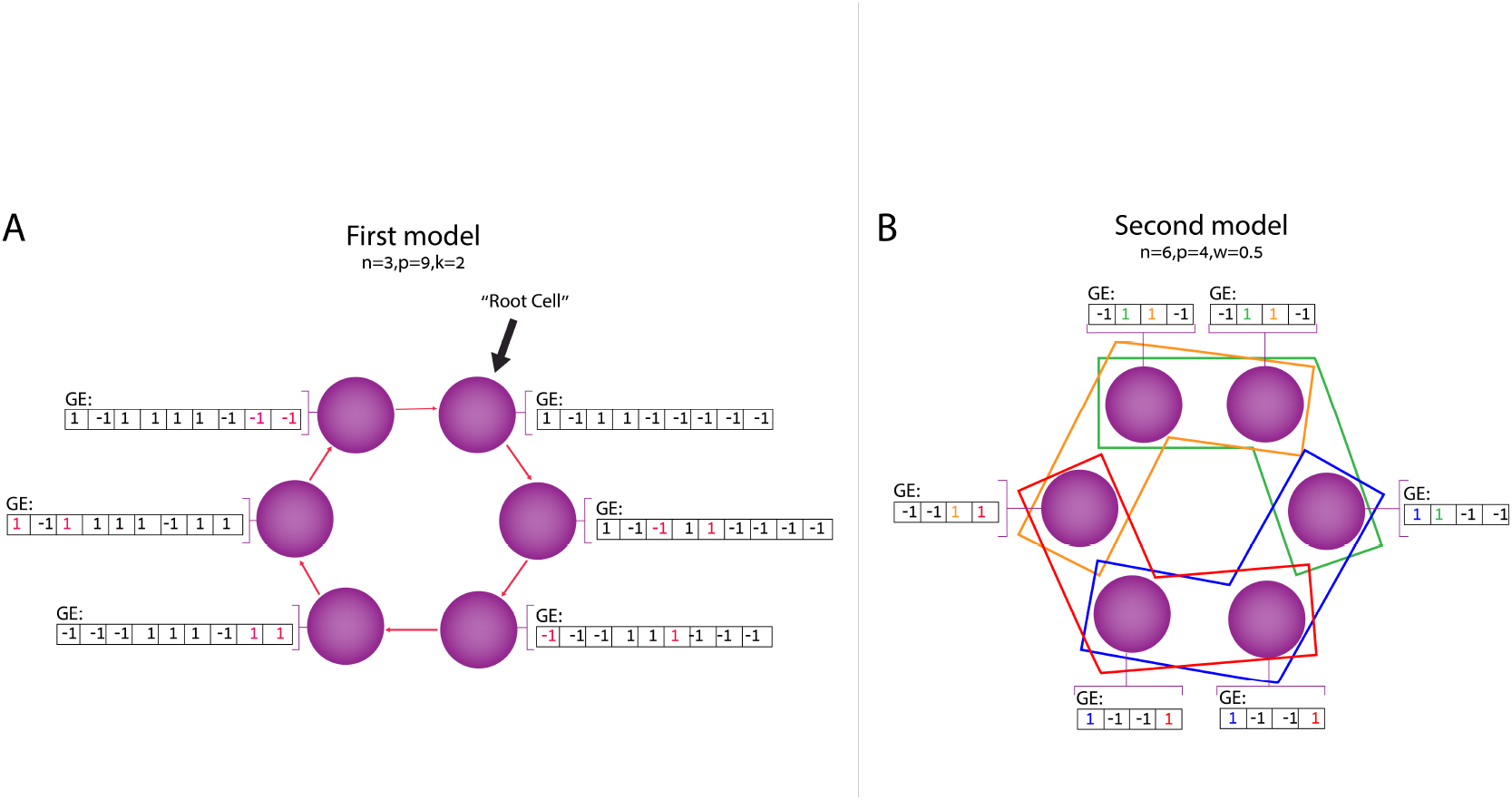
Illustration of the two cyclic signal models: the progression (A) and spatial (B) models.

**Supplementary Figure 2:**
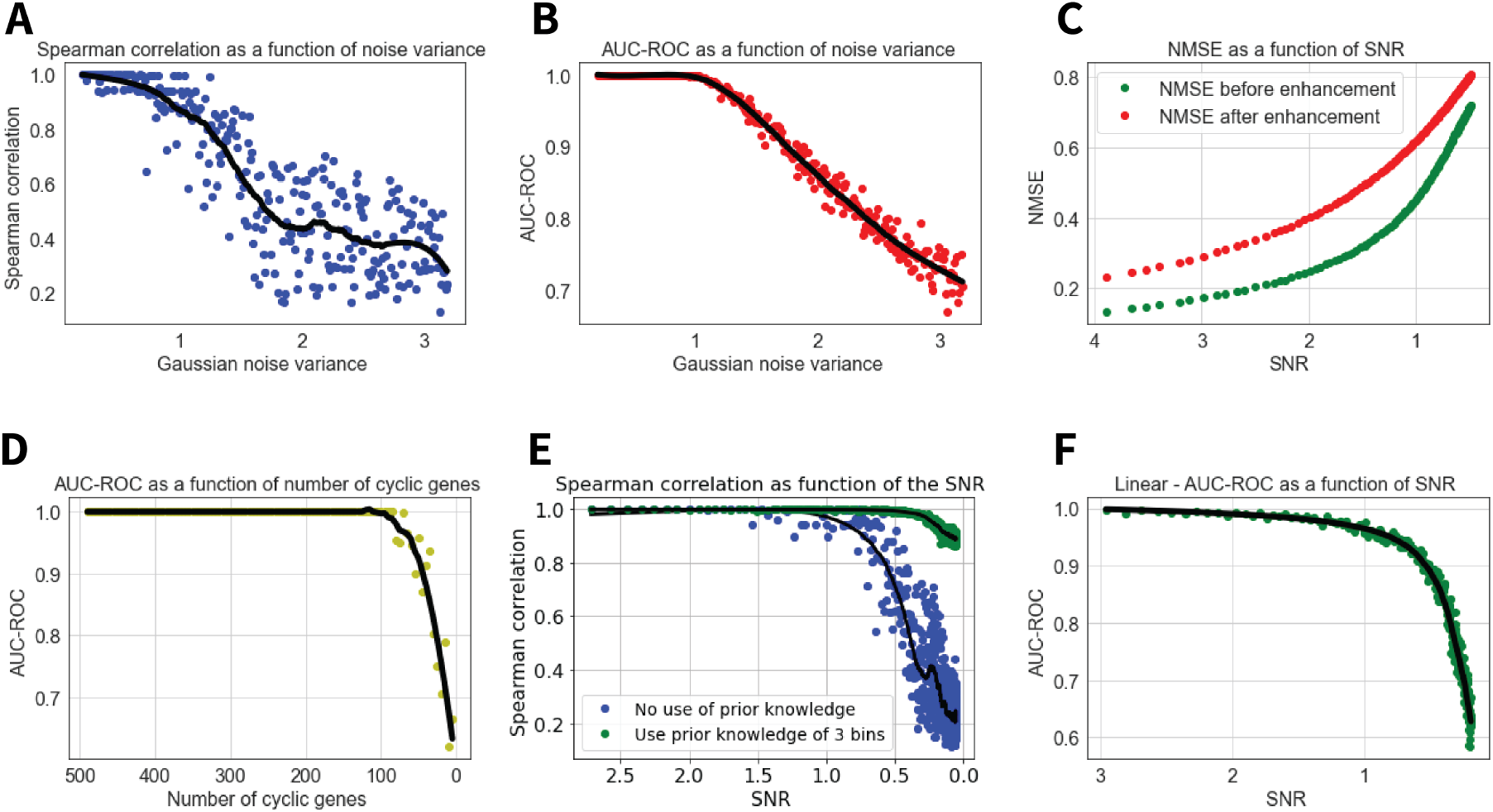
(A) Spearman correlation between the ground-truth permutation and the predicted permutation as a function of noise variance over 300 simulations. The reconstruction algorithm was applied to a simulated gene expression matrix based on a cyclic signal of 100 cells and 500 genes, according to the second model. Gaussian noise was added with varying variance. (B) AUC-ROC of informative genes inference task as a function of noise variance over 300 simulation iterations. A cyclic signal and a lineage signal consisting of 256 cells and 250 genes were simulated in each iteration. These signals were concatenated to form a gene expression matrix of 256 cells and 500 genes, and Gaussian noise with varying variances was added to the matrix entry-wise. (C) NmsE: 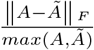 before and after applying the enhancement algorithm over simulated data. (D) Applying the reconstruction algorithm and the genes inference algorithm over simulated data with varying number of cyclic genes. In each simulation, 100 cells were simulated with *x* cyclic genes and 500 – *x* lineage genes. (E) Applying prior knowledge for the reconstruction task. Spearman correlation between the ground-truth permutation and the predicted permutation as a function of SNR. De novo reconstruction (blue), and reconstruction with prior knowledge of low-resolution ordering (3 bins; green), so the mapping of each cell is restricted to the third cycle associated with its label. (F) AUC-ROC of genes inference algorithm as function of noise variance/SNR for inferring linear genes, using the spectrum that is described in methods 4.2.

**Supplementary Figure 3:**
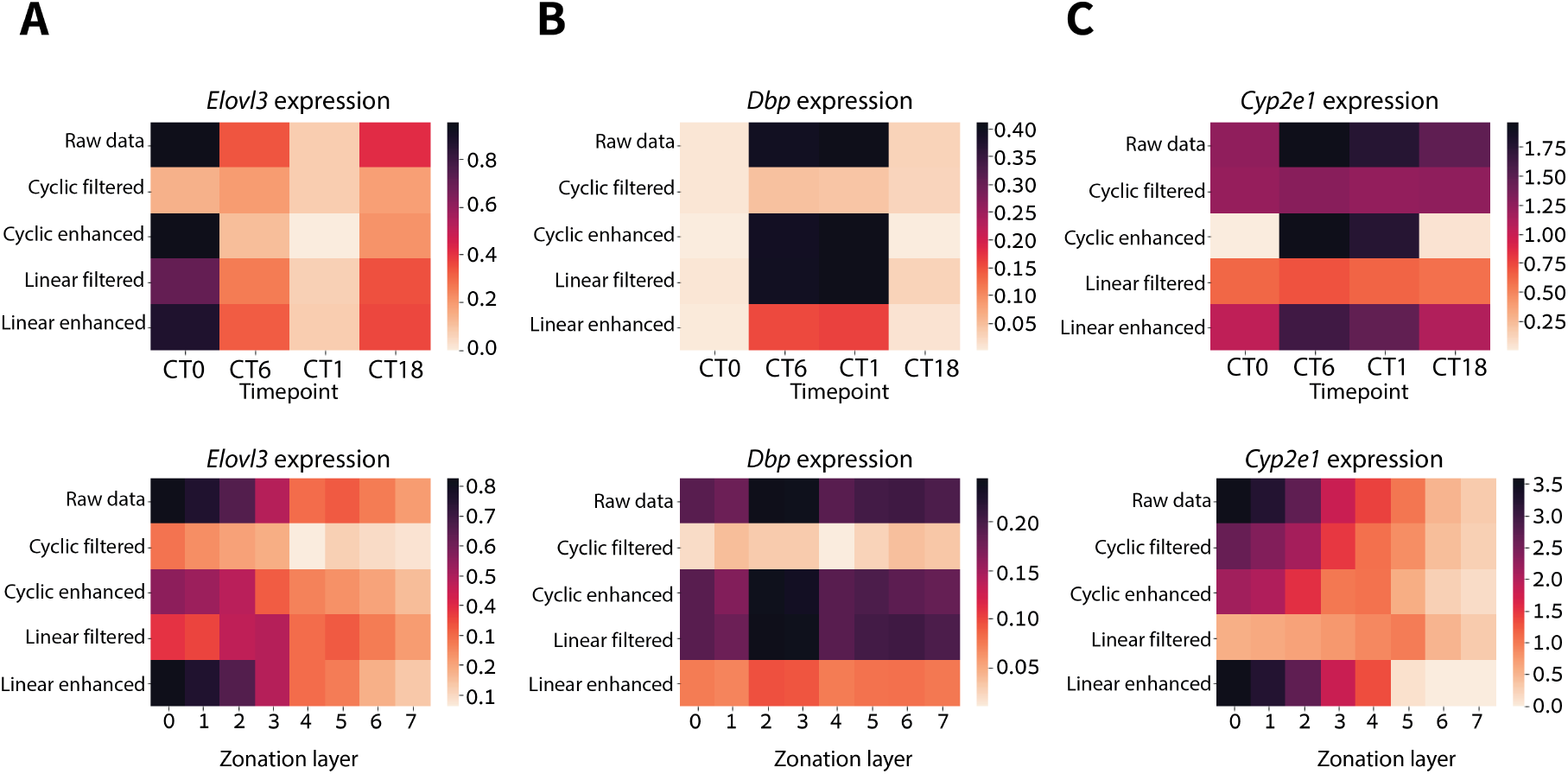
Heatmaps of the expression of *Elovl3* (A), *Dbp* (B) and *Cyp2e1* (C) as a function of the sampling time and the zonation layer.

**Supplementary Figure 4:**
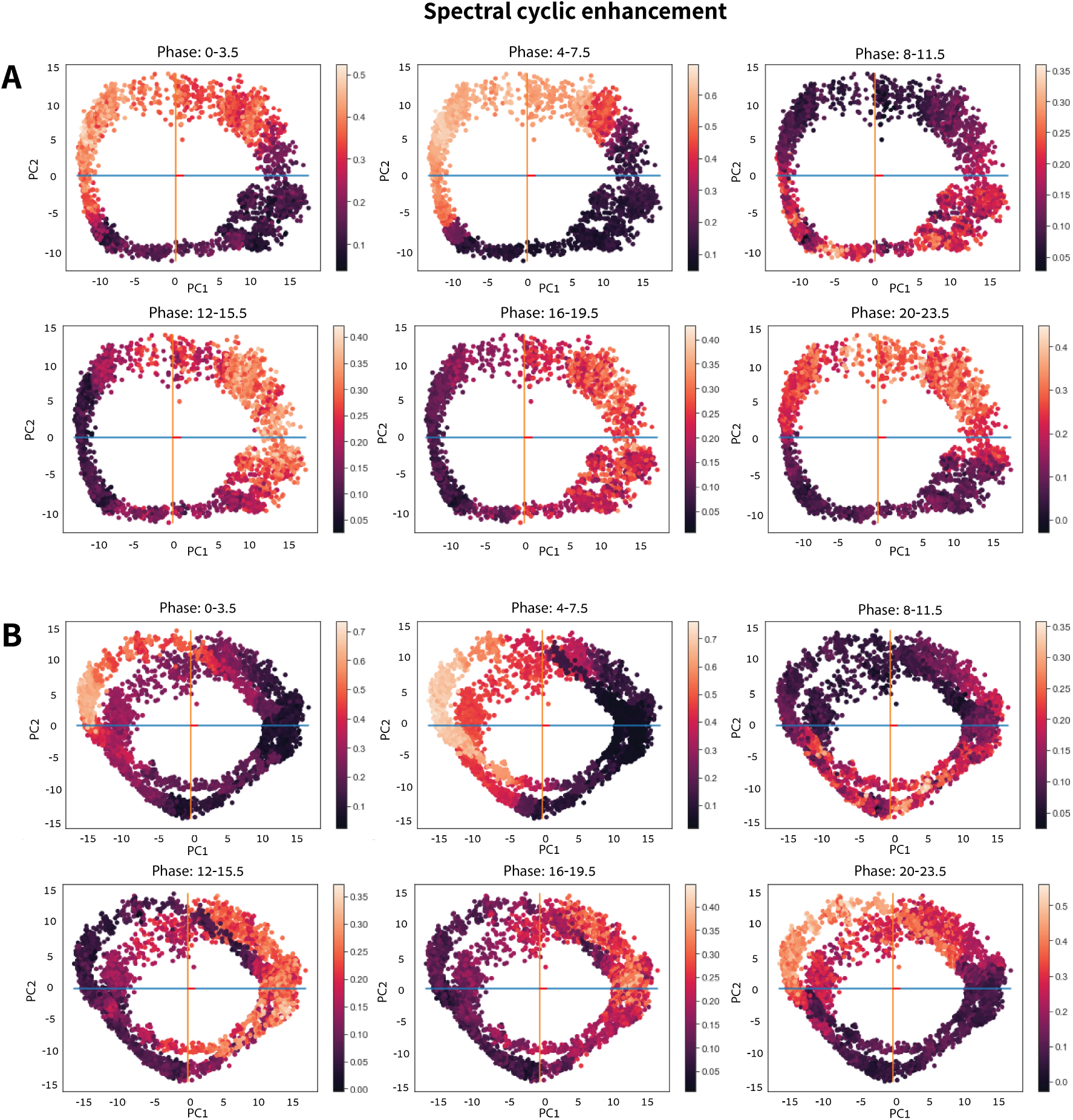
Diurnal cycle in Chlamydomonas. (A) 2 dimensional PCA of enhanced cyclic signal of the iron deficient (Fe-) experiment. (B) 2 dimensional PCA of enhanced cyclic signal of the iron deficient (Fe-) data concatenated with the iron replete (Fe+) data. The 24 hours cycle was divided to 6 segments and the coloring reflects the normalized sum of the marker genes that are associated with each phase. The cyclic signal of each one of the experiments was reconstructed separately.

**Supplementary Figure 5:**
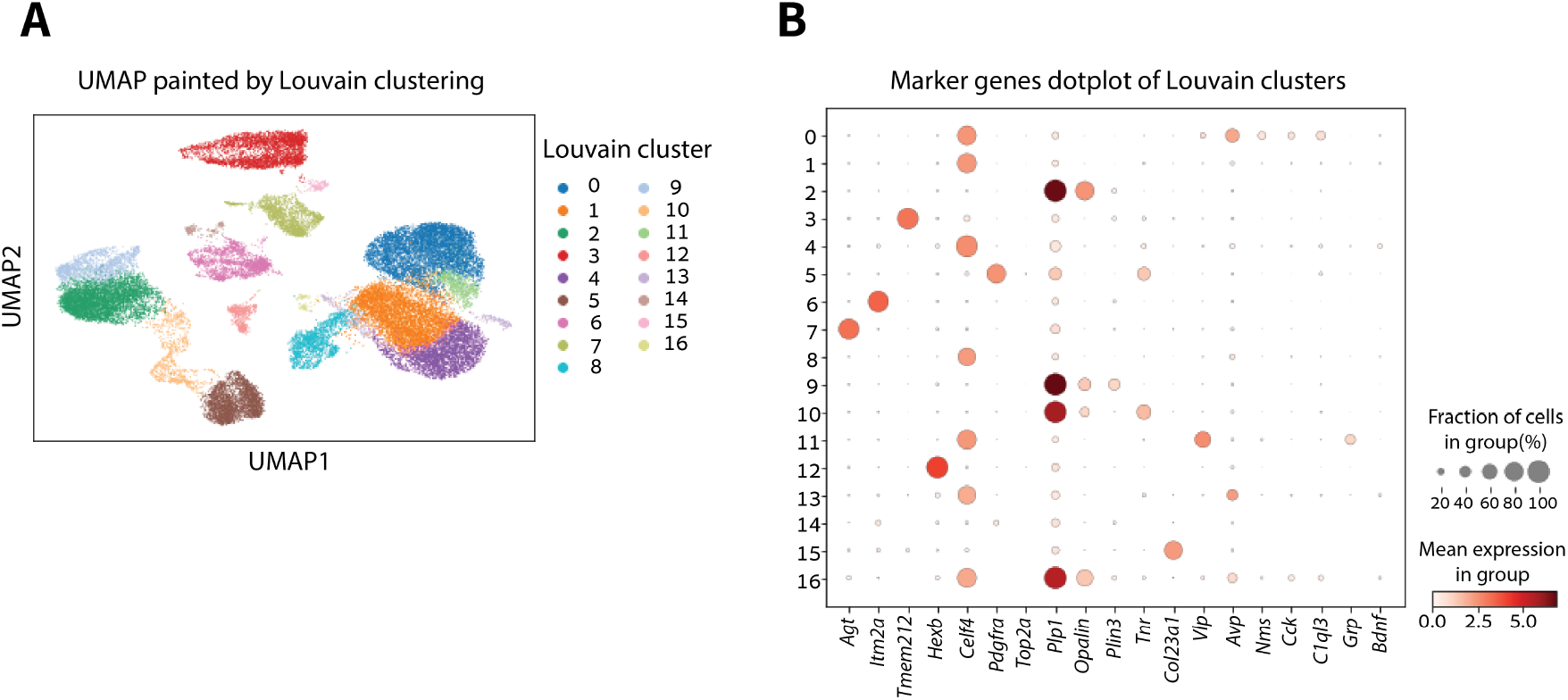
Mice suprachiasmatic nucleus clustering. (A) UMAP of the raw SCN data, colored by clusters that were inferred using the Louvain algorithm. (B) Dotplot of the expression of cell type marker genes in each one of the clusters that were inferred using the Louvain algorithm.

**Supplementary Figure 6:**
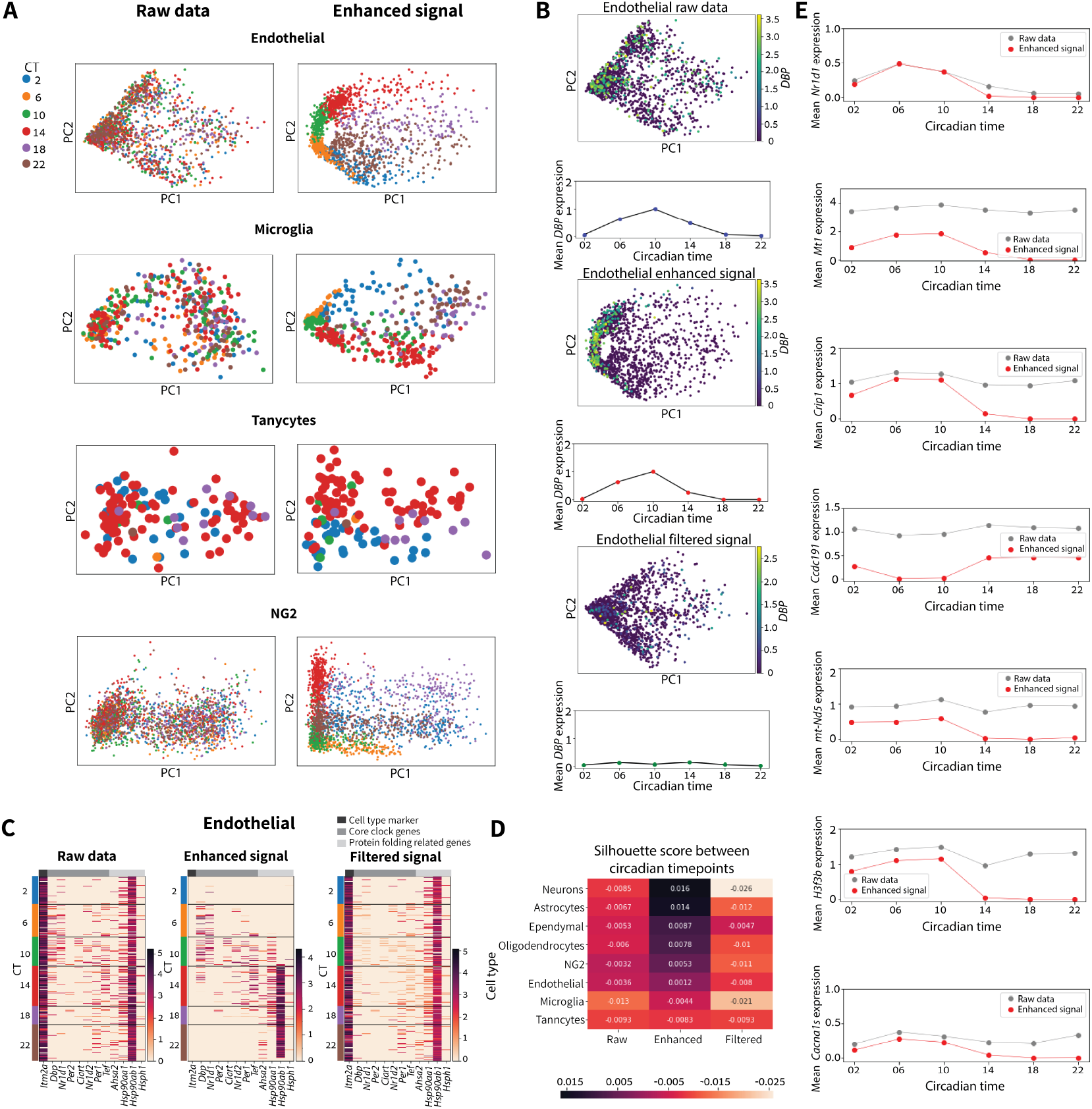
Mice suprachiasmatic nucleus disentanglement. (A) 2D PCA of the raw and enhanced data, colored according to the circadian time for several cell types: endothelial, microglia, tancytes and NG2. (B) Mean expression as a function of circadian time of *Dbp*, 2D PCA colored by *Dbp* expression of raw data, enhanced cyclic signal and filtered cyclic signal for endothelial cells. (C) Heatmaps for raw data, enhanced cyclic signal and filtered cyclic signal of expression of (from right to left) cell type marker genes, rhythmic genes and protein folding genes, for endothelial cells. (D) Silhouette score between circadian timepoints for raw data, enhanced cyclic signal and filtered cyclic signal. Cell types with low fraction of circadian genes (according to [26]) exhibit lower scores. (E) Pre- and postenhancement mean *Nrld1, Mt1, Crip1, Ccdc191, mt-Nd5, H3f3b* and *Cacna1s*. expression as a function of circadian time point. Following cyclic enhancement, regulatory interactions between the transcription factor *Nrld1* and Mtl, *Crip1, Ccdc191, mt-Nd5, H3f3b, Cacna1s* were uncovered.

